# LivecellX: Corrective Deep Learning for Object-Oriented Single-Cell Analysis in Live-Cell Imaging

**DOI:** 10.1101/2025.02.23.639532

**Authors:** Ke Ni, Gaohan Yu, Zhiqian Zheng, Yong Lu, Sophia Hu, Dante Poe, Sijie Zhang, Mia Sanborn, Mostofa Uddin, Zilin Wang, Shiman Zhou, Yanshuo Chen, Xueying Zhan, Weikang Wang, Jianhua Xing

**Affiliations:** Department of Computational and Systems Biology, University of Pittsburgh, Pittsburgh, 15232, PA, USA; Joint CMU-Pitt Ph.D. Program in Computational Biology, University of Pittsburgh, Pittsburgh, 15232, PA, USA; Department of Physics and Astronomy, University of Pittsburgh, Pittsburgh, 15232, PA, USA; Ray and Stephanie Lane Computational Biology Department, Carnegie Mellon University, Pittsburgh, 15213, PA, USA; Department of Computer Science, University of Maryland, College Park, 20742, MD, USA; Institute of Theoretical Physics, Chinese Academy of Sciences, Beijing, 100190, China; School of Physical Sciences, University of Chinese Academy of Sciences, Beijing, 100049, China; UPMC-Hillman Cancer Center, University of Pittsburgh, Pittsburgh, 15232, PA, USA

**Keywords:** live-cell imaging, corrective segmentation network, lineage reconstruction, single-cell trajectory

## Abstract

Live-cell imaging uniquely captures single-cell dynamics in space and time, but robust analysis is limited by segmentation and tracking errors that accumulate across frames. We present LivecellX, a deep-learning–based pipeline that integrates instance-level segmentation error correction with trajectory refinement, leveraging temporal context to recover accurate cell tracks. LivecellX also introduces a benchmark dataset with detailed annotations of common error classes, providing a resource for method development and evaluation. Beyond error correction, the framework incorporates modules for classifying biological processes, reconstructing cell lineages, and analyzing dynamic behaviors. Users can interact with the system programmatically or through a Napari-based graphical interface, enabling flexible integration into diverse workflows. By coupling error-aware correction with comprehensive lineage and dynamics analysis, LivecellX establishes an open, extensible platform that advances the accuracy and scalability of live-cell imaging studies.

## 1 Introduction

Advances in single-cell techniques have transformed the characterization of cellular states-phenotype, transcriptomic, proteomic, epigenomic, and metabolic-and their dynamics. These methods are broadly divided into sequencing- and imaging-based, or alternatively into destructive snapshot approaches and nondestructive methods that enable longitudinal cell tracking. Most sequencing- and multiplex staining-based imaging techniques are destructive [1–8].

Unlike computational inference of cell-state transitions from snapshots, live-cell imaging directly captures spatial and temporal dynamics [9–12], but its adoption lags behind destructive high-throughput methods due to technical challenges. One main challenge is cell segmentation, which must be highly accurate to enable reliable tracking and quantification of features such as morphology and protein distribution [13]. Deep learning, particularly convolutional neural networks and vision transformers, has enabled large-scale segmentation [8, 14–19], and tools like CellPose and *µ*SAM represent current state-of-the-art. However, no method achieves perfect accuracy [20]; segmentation errors, especially under-segmentation(under-seg) and over-segmentation(over-seg), remain common and propagate into tracking, where performance depends heavily on segmentation quality (Fig.1a)[21–25].

Consider a live-cell imaging experiment that images 10^2^–10^4^ cells simultaneously every 10 min for a duration of 16 hours, generating movies with ~ 100 frames. With a segmentation accuracy of 98%, the probability of tracking a trajectory without any segmentation errors is merely approximately 0.98^100^ ≈ 0.13(Fig. 1b). For a seven day experiment, the accuracy further reduces to 0.98^1008^ ≈ 1 × 10^−9^. With a large longtime imaging dataset, e.g., the above-mentioned seven-day experiment with a total of ~ 10^5^–10^7^ cells, it is impractical to identify and correct a fraction of ~ 10^3^–10^5^ missegmented cells through visual inspection. In applications, label-free live-cell imaging emerges as an appealing alternative to fluorescent imaging because of the reduced photo-toxicity and no requirement of genetic engineering [11]. Due to the lack of corresponding training data, segmentation accuracy is typically even lower than that achieved with fluorescent images.

**Fig. 1.**
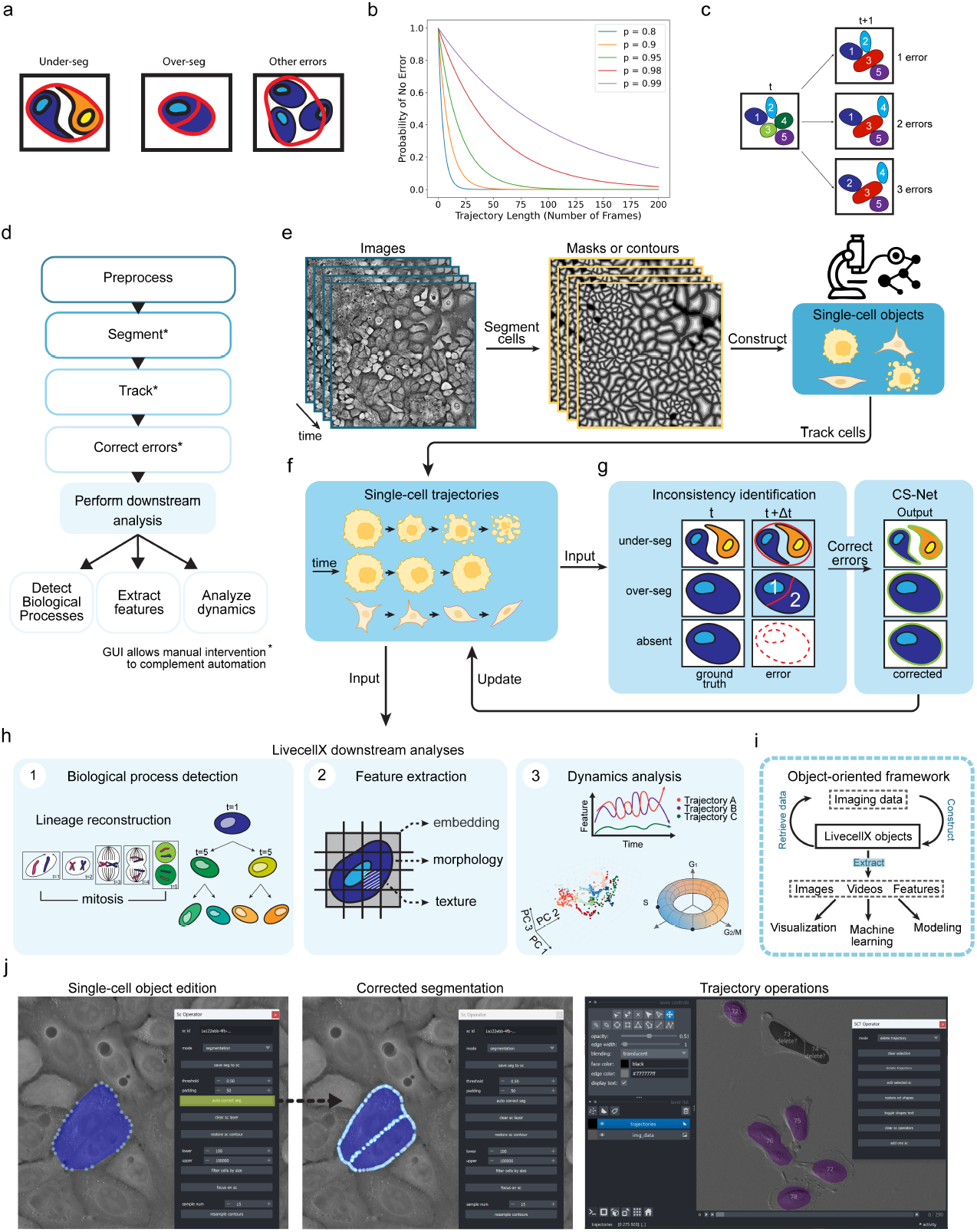
Overview of the LivecellX framework for analyzing single-cell dynamics from live-cell imaging. **a**, Typical segmentation errors in live-cell imaging, including under-segmentation, over-segmentation, and other errors. Ground-truth cell boundaries (black) and predicted contours (red) are shown schematically. **b**, Probability curves illustrating how per-frame segmentation error rates affect the likelihood of obtaining an entirely error-free trajectory as a function of trajectory length. Lines correspond to different per-frame accuracies (80, 90, 95, 98, and 99%), highlighting the rapid decay of error-free probability in long-term imaging even at high accuracy. **c**, Schematic showing how a single segmentation error can propagate into linkage errors in neighboring trajectories. At frame t, cells 3 and 4 are distinct, but at frame t+1 they are merged due to under-seg. Colors indicate ground-truth linkages, while numbers show tracking algorithm predictions. **d**, The LivecellX pipeline integrates segmentation, tracking, error correction, biological process detection, and trajectory-level analysis. **e**, Starting from raw images, cells are segmented into single-cell objects. **f**, Single-cell objects are tracked to form trajectories. **g**, Segmentation errors (under-seg, over-seg, or absent cells) are detected by inconsistencies between consecutive masks and corrected by CS-Net, which takes both the raw image and erroneous mask as input. **h**, LivecellX supports downstream analyses, including lineage reconstruction (left), feature extraction (middle), and dynamics analysis (right). **i**, An object-oriented data framework enables flexible operations in LivecellX. **j**, The GUI supports multiple levels of user interaction, including single-cell object editing (left), segmentation correction (middle), and trajectory operations (right).

Even worse is that segmentation errors often intertwine with single cell trajectory tracking. For instance, one segmentation error in time frame *t* together with movement of cells may generate errors not only in that specific cell trajectory but also in tracking errors in trajectories of neighboring cells (Fig. 1c).

Cell division also increases the complexity of extracting trajectories [26]. Identification of cell division is the key step in lineage tracing that is crucial for analyzing long-term live-cell imaging data [27]. But the scarcity of cell division poses challenges for deep learning models that rely on large and balanced training datasets. For cell dynamics analyses one needs to extract high-dimensional features and choose appropriate representations for cell states [10, 28, 29]. However, limitation in techniques of microscopy and labeling imposes challenges on the widely used multiplex fluorescent labeling for feature extraction in snap-shot imaging experiment. Furthermore, fluorescence labeling can induce significant photo-toxicity in cells during long-term live-cell imaging [27].In recent years, label-free transmitted light imaging emerges as an alternative imaging modality, with significantly enhanced reliability and repeatability of experiments, particularly for long-term live-cell imaging studies.

For label-free or low-plex fluorescent imaging, morphological and textural features are widely used to represent cell states and analyze trajectory dynamics [7, 10, 28, 30– 33]. Hence, software package should include modules for label-free feature extraction and dynamical systems theory-based trajectory analysis[10, 34, 35].

While many computational tools exist for snapshot imaging [7, 8], specialized packages for live-cell imaging remain scarce. Although several solutions support single-cell tracking and trajectory analysis that tackle some but not all the associated challenges simultaneously [26, 36–39]. For instance, most methods rely on stained nuclei [40] and manually curated features, with only a few applying deep learning [26, 41]. The growing scale of live-cell imaging further demands computational efficiency, memory optimization, and dedicated modules for dynamical systems theory-based single cell trajectory analysis.

In this study, we present LivecellX, a comprehensive workflow for analyzing single-cell trajectories from time-lapse imaging datasets. The contribution of LivecellX is threefold. First, it provides an automated procedure to detect and correct under-seg and over-seg errors as well as tracking errors by developing a deep-learning-based method, Corrective Segmentation Network (CS-Net), and integrating temporal information. Second, the LivecellX architecture is designed to transform the analysis task from an image-matrix-oriented processing paradigm to a single-cell object-oriented paradigm. Based on the LivecellX architecture, we integrated methods and designed programming interfaces for tracking, extracting features, detecting biological process, lineage reconstruction, and dynamics analyses. Finally, LivecellX is integrated with Napari to allow interactive user-in-the-loop analyses of live-cell data trajectory objects and cell objects.

## 2 Results

### 2.1 Overview of the LivecellX architecture and workflow

LivecellX is a comprehensive framework for live-cell imaging data analyses (Fig. 1d). It is designed to improve cell segmentation and tracking accuracy based on the additional temporal information between consecutive frames in live-cell imaging, and simplify trajectory dynamics extraction and complex cellular process analysis.

LivecellX takes cell segmentation and trajectory tracking results from other methods as input (Fig. 1e), saves them as a collection of single cell trajectories (Fig. 1f). The segmentation and trajectory results are fed into and iterate between an inconsistency identification module and the CS-Net (Fig. 1g). The identification module exploits the temporal information between consecutive frames for identifying candidates of potentially under-or over-seg cells, and feeds the information to CS-Net. CS-Net is a context-aware, multi-scale machine learning architecture designed to automatically reduce segmentation inaccuracies. It has been pretrained specifically for identifying under-seg and over-seg. With the input from the identification module, CS-Net examines the candidate cells and their neighborhoods for segmentation refinement, then returns to the identification module for reexamination.

After the identification module and CS-Net reach a self-consistent result of high quality trajectories, LivecellX provides three downstream analysis modules. A biological process detection module detects biological processes under interest such as mitosis and apoptosis. Correct detection of mitosis enables reconstruction of cell lineages and downstream analyses (Fig. 1h left). A cell feature extraction module optimized for efficiency via parallelization quantifies high-dimensional features like morphology and texture features and stores the information in single-cell objects (Fig. 1h middle). With a third dynamics analysis module, one represents single cell states and trajectories in a high-dimensional feature space (Fig. 1h right). The quantitative representation enables users to uncover underlying patterns in cell behavior, identify dynamic cellular states, and cluster cells based on temporal representation to reveal processes such as cell cycle progression, response to external stimuli or drug treatments, and disease-related phenotypic changes, etc. All these downstream analyses are implemented based on an object-oriented data framework (Fig. 1i). The framework significantly increases the efficiency of information extraction and manipulation on trajectories.

To ensure robustness and usability, we developed an asynchronous graphical user interface (GUI) based on Napari [42], allowing human-in-the-loop surveillance and intervention at any stage of the analyses. The GUI includes a visualization window that displays microscopy images alongside corresponding segmentation masks and annotations (Fig. 1j). When users encounter errors in specific trajectories—such as segmentation drift or mitotic mislabeling—the GUI enables both visual verification and manual correction (Fig. 1j left and middle, Extended Data Fig. 1a), as well as trajectory-level operations like deletion or refinement (Fig. 1j right,Extended Data Fig. 1b). The LivecellX data structures enable the development of extensible, multipurpose GUIs. For example, two Napari windows can be synchronized and controlled by LivecellX to streamline annotation: when the user clicks ‘annotate,’ both windows automatically advance to the next time frame and highlight the corresponding pair of cell masks. Annotation labels are also easily configurable at launch.

### 2.2 Correction segmentation errors with the CS-Net

Despite recent development of deep-learning-based segmentation methods, segmentation errors are generally unavoidable when processing large imaging datasets. Typical segmentation errors include under-seg where two or more cells are incorrectly segmented as one mask (Fig. 2a top), and over-seg where one cell is incorrectly segmented into two or more masks (Fig. 2a middle) [43–45]. In addition, we defined a subtype of over-seg error termed over-seg dropout error for cases where only a portion of the single-cell is segmented while other portions are not (Fig. 2a bottom).

**Fig. 2.**
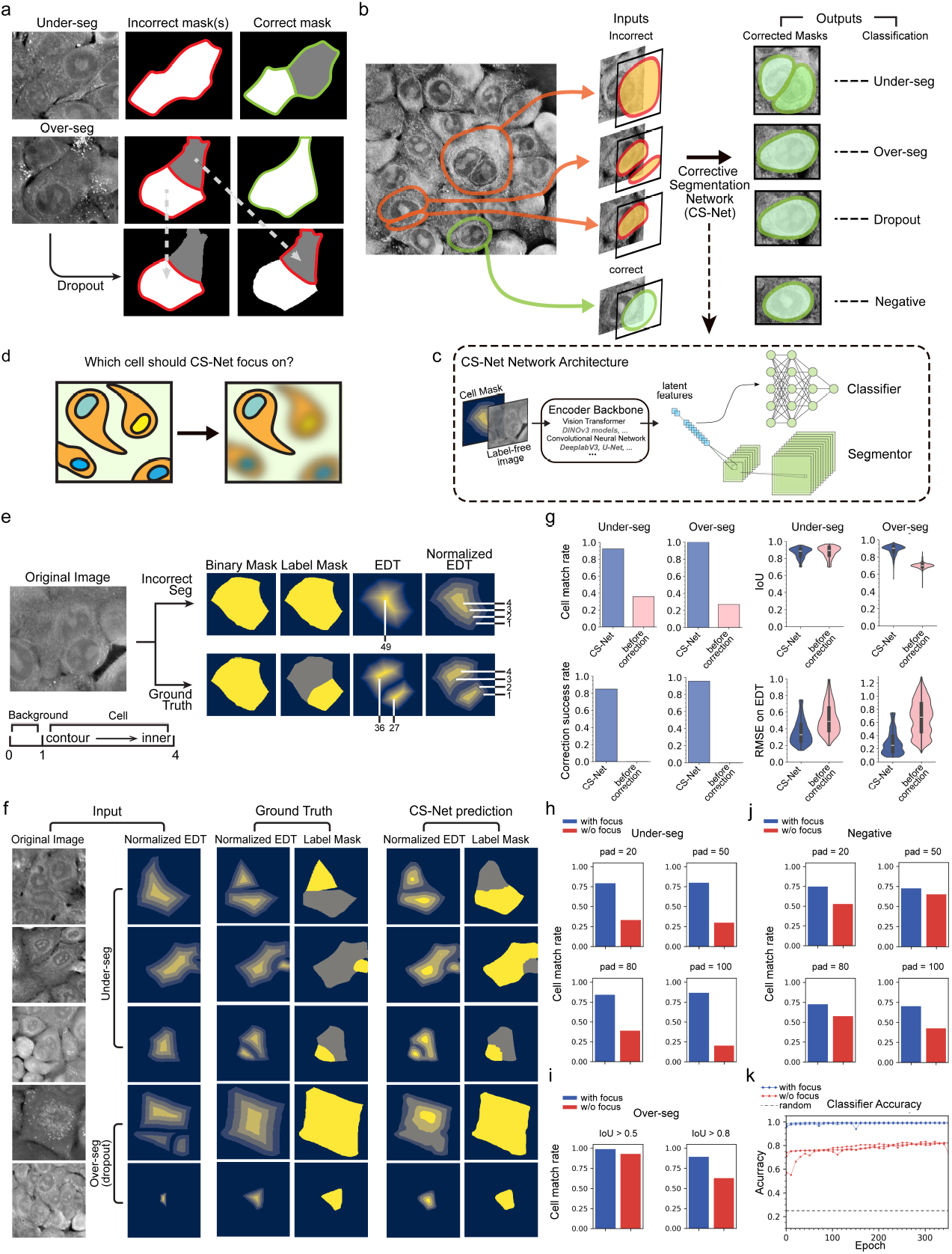
Corrective segmentation and error classification with CS-Net in LivecellX. **a**, Representative examples of segmentation errors in live-cell imaging, including under-seg (top), over-seg (middle), and over-seg dropout cases (bottom). For each, the original image, incorrect mask(s) (red), and correct mask(s) (green) are shown. **b**, Schematic of the CS-Net correction process. Candidate cell masks (incorrect and correct) are extracted from label-free images and input into the CS-Net, which outputs a corrected mask and classifies the error as under-seg, over-seg, dropout, or negative (correct). **c**, CS-Net architecture comprising dual-branch encoders for cell mask and label-free image, followed by segmentation decoder and input-type classifier. **d**, Illustration of the focus mechanism: among adjacent or overlapping cells, the focus mechanism enables CS-Net to correct the designated cell instance. **e**, Different mask types used as the focus mechanism including binary mask, label mask, EDT, and normalized EDT for both ground truth and prediction. **f**, Qualitative examples of CS-Net correction for under-seg, over-seg and over-seg dropout cases. Each row shows the original image, input normalized EDT, ground-truth mask, and CS-Net prediction. **g**, Quantitative evaluation of CS-Net on under-seg and over-seg errors. Metrics shown include correction success rate, cell match rate, IoU, and RMSE of EDT. **h–j**, Cell match rate analysis for under-seg(**h**), over-seg(**i**), and negative (correct)(**j**) cases across varying padding sizes, illustrating consistent performance gains with the focus mechanism; comparison of cell match rates for over-seg at different IoU thresholds (**i**). **k**, Training curves for CS-Net classifier accuracy, demonstrating robust convergence above the random baseline. Classification performance is consistently improved by the focus mechanism.

The candidate segmentation errors are identified using temporal context (see next section) and cropped as region of interest (ROI) from the original images (Fig. 2b). The trained CS-Net takes each ROI– consisting of a candidate cell, its neighborhood, and the segmentation mask –as input and outputs a corrected mask as well as assigns the segmentation status to one of four categories: under-seg, over-seg, over-seg dropout, and no error (negative) (Fig. 2b). An ROI is labeled as negative if the cell is correctly segmented and requires no correction. The CS-Net model comprises a feature encoder—based on either a vision transformer or a convolutional neural network which compresses the ROI into a latent representation, followed by two subnetworks: a segmentor and a classifier (Fig. 2c).

#### 2.2.1 Training of CS-Net

CS-Net needs to be pretrained on curated training and testing datasets independent of inputs from the identification module. A training dataset includes ROI samples of all the four categories mentioned in the above section. To fully leverage annotated data, we implemented synthetic data generation algorithms to introduce both under-seg and over-seg samples, particularly to address class imbalance in our datasets (Extended Data Fig. 2a-e). Having negative samples in the training dataset enhances the CS-Net robustness from altering accurate segmentation results misidentified by the identification module.

A key challenge for CS-Net is to target only erroneous cells within an ROI (Fig. 2d). CS-Net differs from typical segmentation models by concentrating on potentially incorrect cell masks through adding an additional image channel, i.e., a *focus mechanism*. By focusing on individual cell cases, CS-Net allows for precise training and development of specific performance metrics, enhancing control over cell subpopulations. While the focus channel can be the mask or label mask, we found that the Euclidean distance transform (EDT) of the mask (Fig. 2e)[46] work the most robustly on guiding the deep learning model to concentrate during correction (Section 5.2.2). Furthermore, we implemented a normalization technique that scales the EDT values to a fixed range of 1 to 5, producing a contour-map-style representation. This normalized input helps CS-Net prioritize pixels that require correction (Fig. 2e right), and mitigate numerical instability of varying EDT pixel values caused by different cell sizes. The normalized EDT mask and the raw label-free channel image are then jointly fed to CS-Net for correction. During postprocessing, the normalized EDT values further facilitate selecting an appropriate threshold for generating binary masks (Section 5.2.2).

For under-seg, the focus channel highlights regions of individual cells that need separation, whereas over-seg requires identifying multiple fragments of a single cell for merging, which typically relies on manual annotation or auxiliary detection algorithms. To avoid these complexities, we augmented the training data with overseg dropout samples, each containing a single fragment from an over-seg case (see Section 5.2), enabling CS-Net to reconstruct full cell masks from partial, fragmented inputs. Combined with the focus mechanism, CS-Net also employs dataset processing strategies—such as varying ROI padding to include different neighborhood contexts—that enhance generalization and robustness across diverse input sizes and spatial resolutions(Extended Data Fig. 2f).

#### 2.2.2 Evaluating CS-Net

Fig. 2f shows representative corrections of under-seg, over-seg, and over-seg dropout. These inputs span varying scales and background padding, all of which CS-Net robustly handled. The model generalizes across different types of microscopy images with high confluence of cells (Extended Data Fig. 3a). Additionally, CS-Net resolves non-trivial errors beyond canonical under- and over-seg like adjacent-cell merges, boundary ambiguities, three-cell under-seg, and uncategorized mis-segmentation (Extended Data Fig. 3b).

**Fig. 3.**
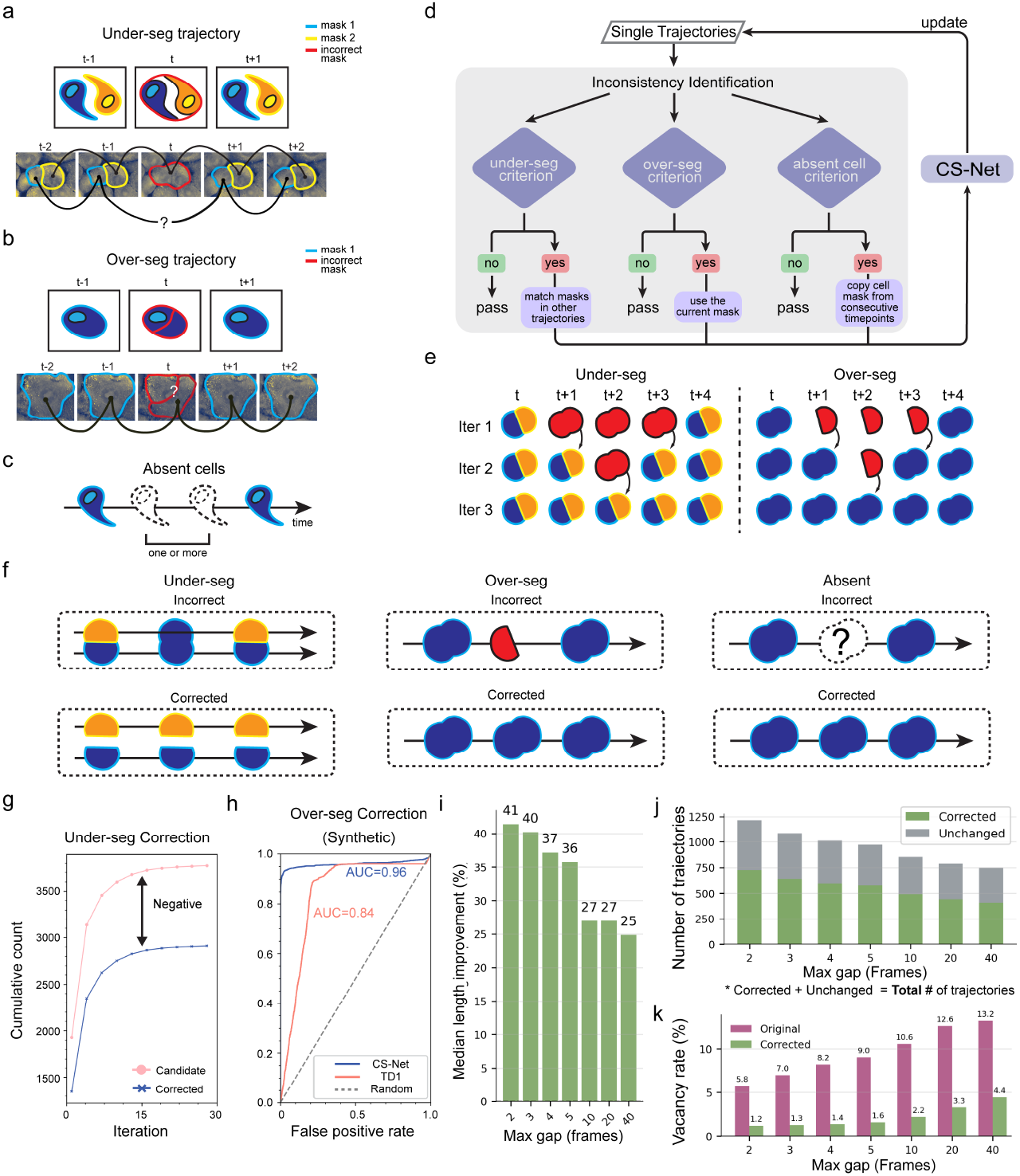
Tracking correction using CS-Net in LivecellX. **a**, Typical tracking error caused by under-seg. Under-seg results in two trajectories merging into one mask at a given time point, causing a missing frame in one trajectory and an incorrect cell mask at this frame in the other. **b**, Typical tracking error caused by over-seg. A trajectory is split at a single time point, leading to sharp morphological changes and short-lived tracklets due fragmented masks. **c**, Absent cells result in missing frames in the trajectory. **d**, Flowchart of the trajectory correction pipeline. Each trajectory is screened for temporal inconsistencies. Candidate frames are classified as under-seg, over-seg, or absent cell using dedicated criteria, and then corrected by CS-Net. Corrected masks are iteratively updated within the trajectory. **e**, Illustration of multi-iteration correction for persistent under-seg (left) and over-seg (right) errors across several consecutive frames. **f**, Examples of linkage updates in single-cell trajectories before (incorrect) and after (corrected) segmentation error correction for under-seg (left), over-seg (middle), and absent cell errors (right). Arrow represents time flow. **g**, Cumulative counts of under-seg candidates and successfully corrected cases over multiple correction rounds. The difference reflects negative candidates identified by temporal inconsistency screening. Note that TD-1 is designed to have potential negative candidates for subsequent CS-Net identification. This expanded list minimizes the occurrence of false-negative cases. **h**, Receiver operating characteristic (ROC) analysis for over-seg correction, comparing CS-Net correction results to a simple feature change score. CS-Net achieves superior discrimination (AUC = 0.96). **i**, Median trajectory length improvement (%) after applying correction, plotted against the maximum gap size (frames) in trajectory linking. **j**, Number of trajectories classified as corrected (green) or unchanged (gray) for each maximum gap size; each bar represents the total number of trajectories, not only candidate cases. **k**, Vacancy rate (%) of trajectories before (gray) and after (green) correction, shown across different maximum gap sizes.

We evaluated segmentation performance using four metrics: cell match rate, correction success rate, intersection over union (IoU) calculated on cell contours (morphology), and root mean square error (RMSE) of EDT masks (Extended Data Fig. 3c-e; see Methods for definitions). Our evaluation employs instance-level mask-based metrics that are more sensitive to mask errors.

Statistical analysis of mask-based metrics showed that CS-Net with the focus mechanism significantly improved segmentation quality (Fig. 2g). CS-Net successfully corrected nearly all over-seg error cases. In under-seg scenarios, CS-Net achieved strong performance across three of the four evaluation metrics, with the exception of the conventional binary mask IoU, which is commonly used in the field (Fig. 2g). This metric is not well suited for evaluating under-seg, as erroneous and ground-truth masks often appear nearly identical at the pixel level despite representing different cell instances(Fig. 2f). Consequently, even a perfect correction by CS-Net may result in minimal or no change in the binary mask IoU. In contrast, RMSE of EDT masks, cell match rate, and correction success rate provide more sensitive and informative assessments for under-seg cases by directly evaluating instance correspondence and morphological accuracy.

We compared the performance of CS-Net with and without the focus mechanism under different conditions (Fig. 2h–j). For under-seg and negative cases, the performance gap between the two models increases as padding size grows, reflecting the presence of more cells in the label-free image crop. For over-seg cases, the gap widens as the IoU threshold becomes more stringent, demonstrating the advantage of the focus mechanism under stricter criteria. When no padding is applied, however, overseg crops often contain only a single cell, in which case the focus mechanism provides little benefit. Overall, CS-Net with the focus mechanism achieved consistently higher cell match percentages across all conditions, maintaining robust performance across varying background sizes, as controlled by different padding values from raw imaging data (Extend Data Fig. 3 f-h). Notably, the performance gap favoring the focus mechanism expanded with larger padding sizes, underscoring its effectiveness in handling diverse spatial contexts. The comparison of correction success rates at different padding size, IoU thresholds, microscopy modalities and error types on in different data subsets also reveals the robustness of this method(Extend Data Fig. 3 i-j).

The accuracy of the classifier was also higher when CS-Net is equipped with the focus mechanism. This aligns with intuition: without the focus mechanism, CS-Net lacks information about which specific cell within the ROI should be classified and segmented, leading to reduced performances (Fig. 2k).

Across encoder backbones, we observed no consistent advantage of ViT-based over convolutional neural network (CNN)-based backbones when parameter counts were comparable. Instead, scale dominated: larger encoders yielded better corrections. Within DINOv3 backbones, a fine-tuned large variant (ViT-7B/16) substantially out-performed fine-tuned small variants (e.g., ViT-S/16): correction success rate with small ViT models could drop from ~ 0.7 to ~ 0.3 on our benchmarks, whereas the larger model maintained high success. A similar scale effect held for CNN backbones, with larger models consistently outperforming smaller ones under matched training settings (see Methods for configurations).

### 2.3 Linkage correction in single cell trajectories with CS-Net

Cells are deformable, often exist in high-density environments, and can divide, disappear, or occlude each other. While existing tracking algorithms have been adapted to account for these biological constraints, they generally assume that the input segmentation masks are accurate [47–49].

However, segmentation errors cause several typical linkage distortions between single-cell masks including cascade of wrong linkages involving neighboring cells caused by under-seg of two cells from different trajectories(Fig. 3a), decreased trajectory quality and erroneous linkages caused by unassigned over-seg fragments (Fig. 3b), missing frames due to absent cells (Fig. 3c, Extended Data Fig. 4a, details in Methods). Linkage errors can be further complicated when more neighboring cells are involved.

**Fig. 4.**
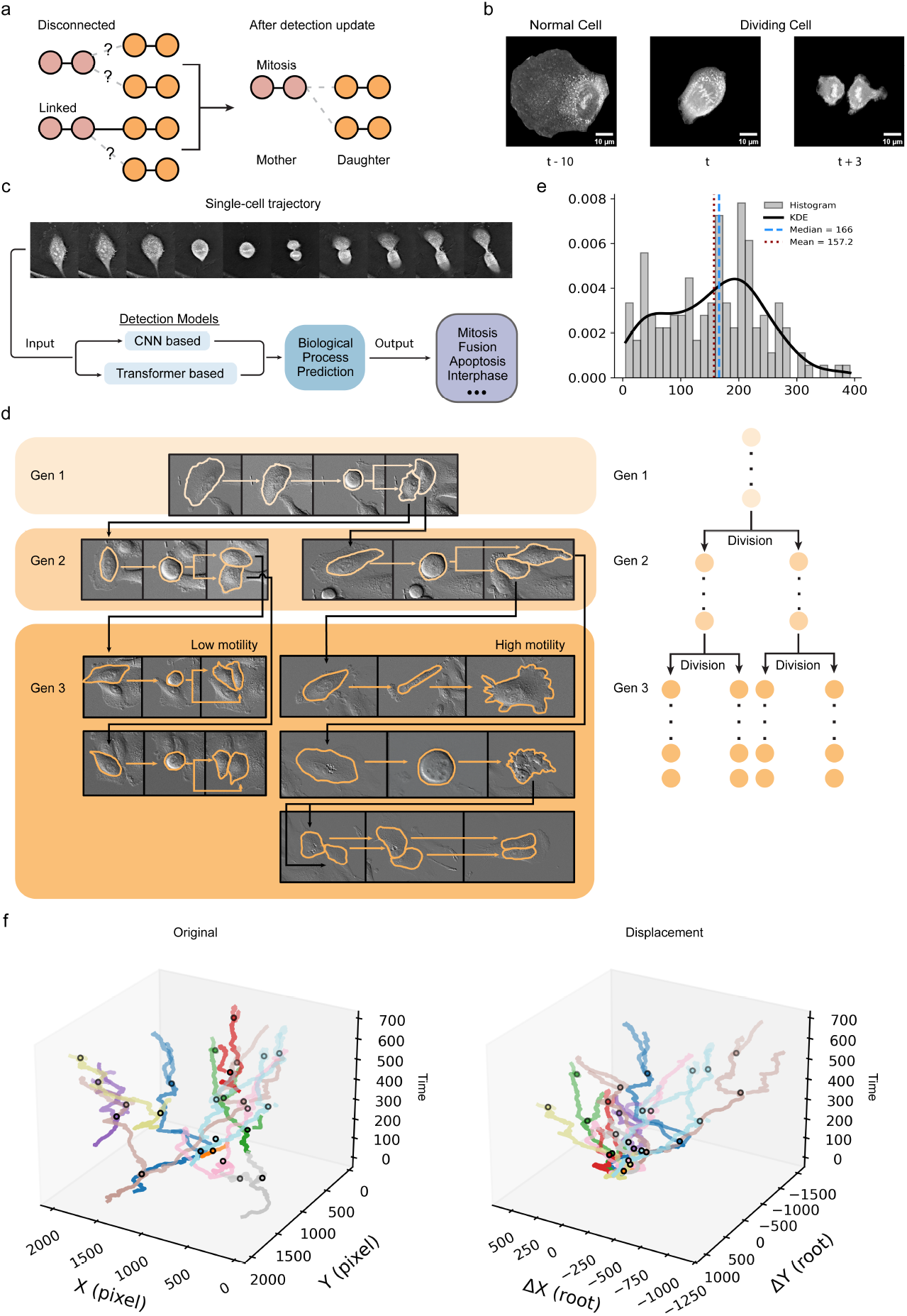
Automated cell lineage reconstruction and biological process detection in Live-cellX. **a**, Solution of linkage errors caused by mitosis events. In standard tracking, mitosis can lead to disconnected or ambiguous associations between mother and daughter cells. LivecellX detects mitosis events and accurately establishes lineage linkages between mother and both daughter cells. **b**, Distinctive appearances of cells before and after cell division in one single cell trajectory. **c**, Schematic of the biological process detection module. Single-cell trajectories are analyzed by CNN-based or transformer-based models to predict mitosis and other biological events, enabling automated lineage tree construction. **d**, Representative reconstructed cell lineage across three generations, visualized with cell contours in microscopy images. Generation 3 cells are further categorized by low and high motility, highlighting heterogeneity in movement within lineages. **e**, Distribution of cell cycle durations for reconstructed lineages, shown as a histogram with kernel density estimation (KDE). Mean and median durations are indicated. **f**, Three-dimensional visualization of cell lineage trajectories over time, shown as absolute coordinates (left) and as displacements from the root cell (right), revealing both lineage structure and motility patterns.

Robust lineage reconstruction requires jointly correcting segmentation masks and tracking linkages, motivating integrated approaches that iteratively refine both. In CS-Net, candidate errors are first detected and cropped as ROIs. Note that there are typically strong overlap between correct masks across consecutive frames, and segmentation errors appear as abrupt changes in overlap or morphology (Fig. 3a-c). To capture these, an inconsistency detection module (TD-1) filters temporal inconsistencies using IoU and Intersection over Minimum (IoMin) metrics (Extended Data Fig. 4b). Each under-seg candidate was represented as a triplet 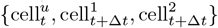 where cell 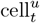 denotes the under-seg mask. Each over-seg candidate was represented as a pair 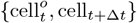, where cell 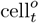 denotes the over-seg mask (Extended Data Fig. 4c). Candidates meeting the error criteria are cropped with surrounding regions and passed to CS-Net for correction (Fig. 3d). The candidate triplets are affected by the threshold values of IoU and IoMin scores (Extended Data Fig. 4d-e). In addition, the efficiency of detection is affected by situations like mitosis and cells with high motility.

Testing on multiple datasets revealed that the TD-1 method alone was insufficient for detecting segmentation errors, particularly when errors occurred in consecutive frames along a single-cell trajectory (Extended Data Fig. 4f). To address this limitation, we developed a multi-iteration, trajectory-level algorithm that employs CS-Net to iteratively correct both segmentation and track linkage errors (Fig. 3e). In each iteration, trajectory-level errors are identified, corrected by CS-Net, and the corresponding trajectories updated. This process ensures that the first error in a sequence of persistent errors is resolved in the first iteration, while subsequent errors are corrected in later iterations. Typical linkage errors—caused by under-seg (Fig. 3f, left), over-seg (Fig. 3f, middle), and absent cells (Fig. 3f, right)—are successfully corrected by this approach.

Validation on A549 trajectories showed that, after fifty iterations, the number of attempted and resolved under-seg corrections reached saturation (Fig. 3g). As expected, a clear gap remained between potential errors flagged by TD-1 and true errors corrected by CS-Net, reflecting that CS-Net can detect false positives included in TD-1 detection (Extended Data Fig. 4g-h). The ROC analysis (Fig. 3h) further demonstrated that CS-Net with trajectory inconsistency detection achieved an AUC of 97%, outperforming the 87% AUC from TD-1 features alone. Visual inspection confirmed that all true over-seg errors were correctly identified and corrected.

Next, we compared trajectory lengths before and after our correction procedure (Fig. 3i). For each corrected trajectory, we calculated the relative increase in length. Across all tested settings of the maximum age parameter(*max gap*)in the tracking algorithm (Methods), the median relative increase exceeded 25%, demonstrating the robustness of our correction method (Fig. 3i). Notably, when max gap was small, the relative increase approached 40%, as shorter *max gap* values tend to produce more fragmented trajectories. In line with this, the total number of trajectories decreased substantially after correction, reflecting the successful linking of short trajectory fragments (Fig. 3j).

To further assess performance, we quantified the trajectory vacancy rate, defined as the proportion of missing frames due to factors such as segmentation errors or cells moving out of the field of view. Approximately half of these vacancies were resolved following correction, regardless of the *max gap* setting, underscoring the method’s effectiveness in improving trajectory continuity and quality (Fig. 3k).

### 2.4 A biological process detection module allows user-specified cell biology event-aware image analyses

Tracking cells in live-cell images is uniquely challenging due to biological processes such as division and apoptosis. Cell division, in particular, adds complexity to single-cell dynamics and lineage analysis. When a mother cell divides, its trajectory should terminate while two daughter trajectories are initiated, with the mother–daughter relationship preserved for lineage reconstruction. Without accurate mitosis detection, tracking algorithms may incorrectly link trajectories or miss lineage relationships (Fig. 4a).

Therefore, we developed a module to identify frames where user-specified biological processes occur. As an example, we trained a network to detect mitosis, which is marked by rapid morphological changes; in 2D cultures, both mother and daughter cells appear small and round (Fig. 4b).

We integrated video-based deep learning methods by leveraging full temporal and visual context. Two frameworks — CNN- and transformer-based (Fig. 4c) — were fine-tuned from MMAction video-classification models [50] and adapted to treat mitosis detection as a single-cell trajectory classification task. Detected events are stored in the LivecellX trajectory structure, though manual annotation may still be required for data from different microscopes or conditions. Alternatively, users can identify divisions via the GUI.

Fig. 4d shows an A549 single-cell lineage spanning three generations, revealing differences in motility and cell cycle duration among sister and cousin cells. LivecellX also enables statistical analysis of cell cycle durations (Fig. 4e) and visualization of multiple reconstructed lineages across generations, including spatial positions (Fig. 4f, left) and displacements relative to the initial frame (Fig. 4f, right).

### 2.5 Feature extraction and dynamics analysis modules provide tools for cell dynamics studies

A key downstream task in LivecellX is extracting multiplex features from fluorescent and/or label-free images to represent single-cell trajectories in multidimensional feature space. These features enable visualization, interpretation, and modeling of cellular dynamics. LivecellX integrates tools such as `scikit-image` and `mahotas` to extract morphological and texture-based descriptors (e.g., Haralick features), which strongly correlate with cell states and capture dynamic changes over time. Importantly, many morphological features can be quantified from transmitted-light images, avoiding the limitations of fluorescence labeling, including restricted marker capacity and phototoxicity.

As a demonstration, we extracted multidimensional morphological and texture features from time-lapse images of two cell lines, MCF10A and A549. Feature distributions differed between the lines (Fig. 5a), and cell line–specific clusters emerged in t-SNE and UMAP embeddings (Fig. 5b). These embeddings correlated with specific morphological and texture features(Fig. 5c, Extend Data Fig. 5a), whose contributions were further examined by principal component analysis (PCA) (Extended Data Fig. 5b).

**Fig. 5.**
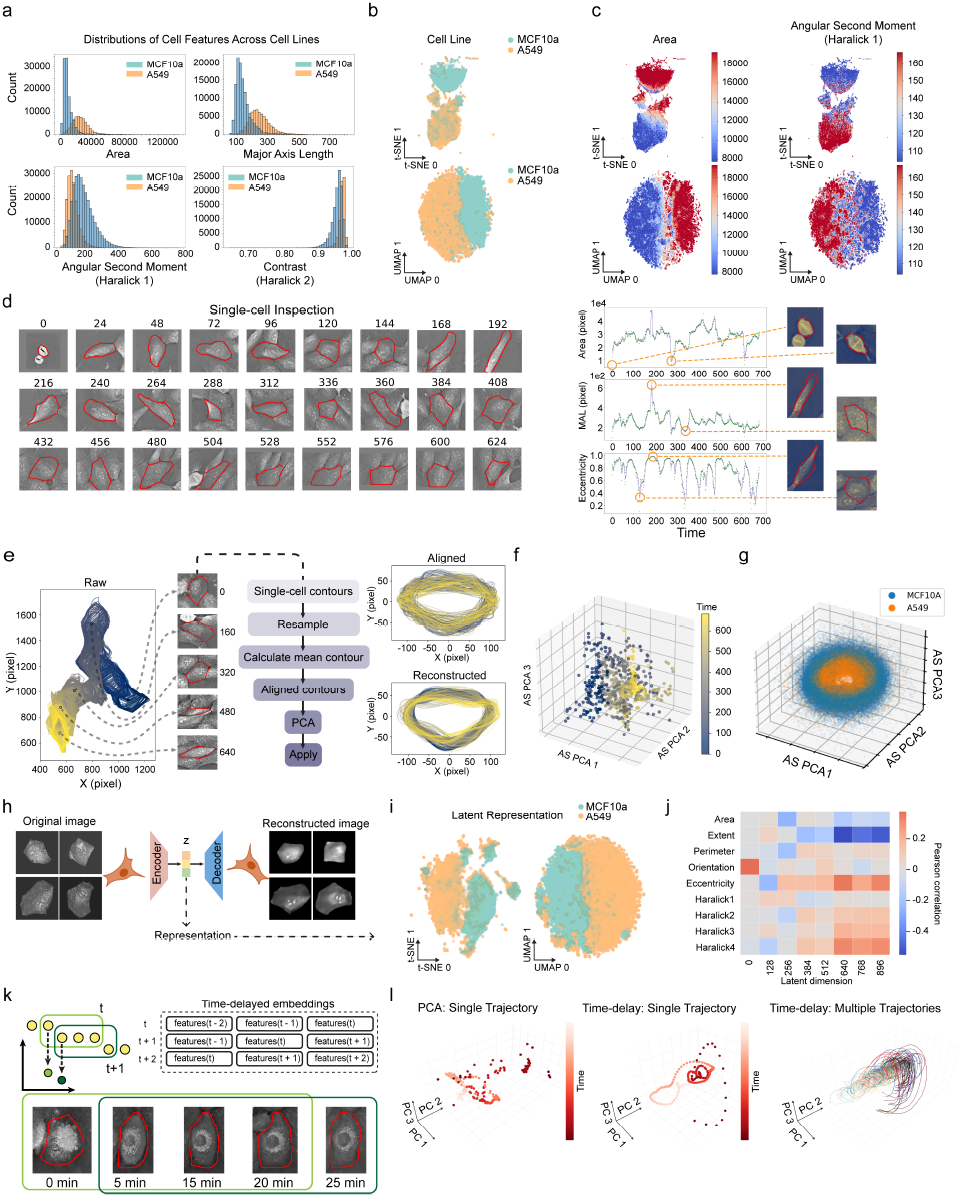
Automated extraction and analysis of single-cell dynamics in LivecellX. **a**, Distributions of morphological (area, major axis length) and Haralick texture features (angular second moment, contrast) of A549 and MCF10A cells. **b** Projections of single-cell morphology and texture features using t-SNE (top) and UMAP (bottom). Colors represent cell lines. **c**, Projections of single-cell morphology and texture features using t-SNE (top) and UMAP (bottom). Colors represent area (left) and angular second moment(right). **d**, Temporal variation of morphological features (area, MAL, and eccentricity) along a single A549 cell trajectory. Left: sequential cell images and outlines. Right: feature dynamics and cell images at extreme feature values. **e**, Pipeline of AS model for quantifying dynamic cell contours. Left: single-cell contours colored by time. Middle: schematic workflow of AS model. Right: aligned (top) and PCA-reconstructed (bottom) contours of a single trajectory. **f**, Trajectory dynamics projected onto the first three PCs of the AS model, colored by time. **g**, PCA projection of AS model representations for A549 and MCF10A cells, revealing cell line–specific characteristics. **h**, Schematic of the Harmony deep learning model for extracting latent embeddings from single-cell images, with example reconstructed images. **i**, Visualization of Harmony latent representations with t-SNE and UMAP, colored by cell line. **j**, Correlations between Harmony latent features and traditional morphological or texture features, displayed as a heatmap. **k**, Illustration of time-delayed embedding applied to single-cell trajectories. Example cell images and masks (red outlines) are shown at different time points. **l**, Time delay embedding of cell cycle trajectory. Left: One cell cycle trajectory in the Harmony embedding feature space. Middle: Time delay embedding of the trajectory on the left panel. Right: Multiple time-delay embedding cell cycle trajectories. PCA is applied for visualization.

LivecellX enhances the quality of extracted trajectories in long-term live cell imaging and enables exploration of dynamic changes along each trajectory. The original images at different time points of a typical single cell trajectory can be retrieved conveniently with data framework of LivecellX (Fig. 5d left). The morphology features including cell area, major axis length(MAL) and eccentricity are shown as time series and can be analyzed in further study(Fig. 5d right). The framework also supports intuitive selection and inspection of individual cells across time points.

In addition to morphology and texture features, LivecellX integrates an active shape (AS) model to represent cell contours [32](Fig. 5e). For example, a representative MCF10A cell trajectory is shown in pixel space with time-colored morphology colored (Fig. 5e left). After alignment the contours are represented with the AS model which extracts multi-dimensional features from single cell contours(Fig. 5e middle), and PCA yields orthogonal modes that describe and reconstruct cell shapes (Fig. 5e right, Extend Data Fig. 5c). Single cell trajectories can then be represented in PCA coordinates (Fig. 5f), capturing dynamic changes in morphology [10, 32]. Applying this model across all trajectories reveals distinct shape dynamics between A549 and MCF10A cells (Fig. 5g).

To exploit the power of machine learning for capturing richer image features beyond handcrafted feature-based methods, we integrated a machine-learning–based feature extractor, Harmony, a variational autoencoder (VAE) designed to disentangle semantic content from transformations and yield invariant, interpretable embeddings [51] (Fig. 5h).

Each cell was represented by a 512-dimensional Harmony embedding, which, when projected using UMAP or t-SNE, yielded clusters comparable to those obtained from traditional morphological and texture features(Fig. 5i, Extended Data Fig. 5d). However, Harmony embeddings contained more uncorrelated features, likely due to their higher information content and the non-linear encoder structure (Extended Data Fig. 5e). To interpret these latent representations, we correlated Harmony embeddings with morphological descriptors, finding strong associations—particularly between the standard deviation component of Harmony embeddings and several morphological and Haralick texture features (Fig. 5j, Extended Data Fig. 5f)

Cellular dynamics can be studied through large collections of single-cell trajectories by reconstructing the underlying dynamic manifold via time-delay embedding, as guaranteed by Takens’ theorem [52]. For a cell at time *t*, this embedding concatenates feature vectors over the range (*t* − Δ*t,t* + Δ*t*) and can be built from morphology, texture, ASM, or VAE embeddings (Fig. 5k). Using Harmony features, time-delay embedding of a cell cycle trajectory revealed the expected cyclic structure (Fig. 5l).

### 2.6 LivecellX is a portable single-cell object-oriented framework with Napari interface

Current image-centric live-cell imaging workflows are computationally inefficient, particularly with heterogeneous formats and large datasets, as they rely on matrix-based operations that complicate object tracking. LivecellX overcomes these challenges with an object-oriented, cell-centric design that organizes data around single-cell objects, assembled post-segmentation and integrated into trajectories for efficient analysis (Fig. 6). The framework operates across three levels: (1) Single-cell objects, encapsulating features, metadata, and subcellular details with interfaces compatible with tools like Cellpose and StarDist [8, 53, 54]; (2) Single-cell trajectories, linking objects over time with metadata such as velocity and lineage for automated reconstruction; and (3) Trajectory collections, which organize all trajectories for large-scale statistical analyses and cross-dataset integration. The emphasis on cell-centric data processing reduces repetitive computations and enhances data interpretability across diverse experimental conditions, enabling large-scale, multi-dataset integration.

**Fig. 6.**
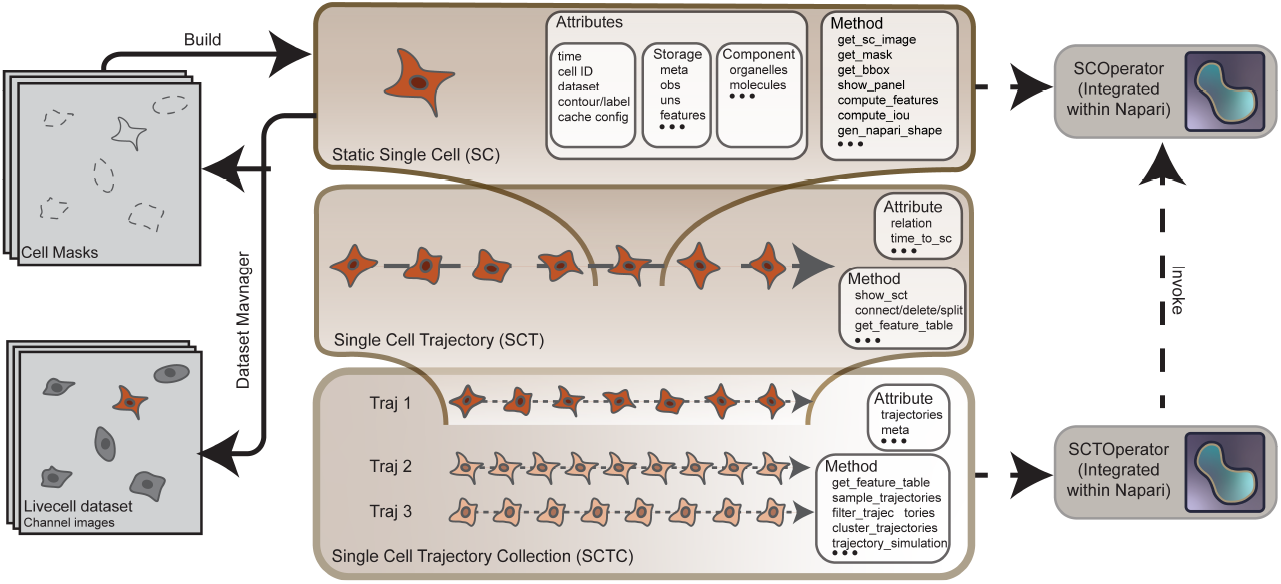
Object-oriented architecture and user interface integration of LivecellX. Hierarchical data structures for representing static single cells (SC), single-cell trajectories (SCT), and collections of trajectories (SCTC). Key attributes and methods enable flexible querying, feature computation, and Napari-based interaction via SCOperator and SCTOperator modules.

A Napari-based GUI leverages the above structure to provide interactive, programmatic access for inspecting, editing, and correcting results, particularly at the trajectory level (Extended Data Fig. 6a–b). LivecellX further optimizes data management through dynamic structures and caching mechanisms (e.g., LRU cache), reducing redundant computation and disk access (Extended Data Fig. 6c).

We integrated LivecellX with PyTorch and different machine learning method to enable on-the-fly data generation and stochastic augmentation from single-cell or trajectory objects (Extended Data Fig. 6c). For instance, a diffusion model is trained for generating single cell images and related masks for training as well as other analyses(Extended Data Fig. 6d).

## 3 Discussion

In this study, we present LivecellX, a comprehensive pipeline for analyzing single-cell dynamics from time-lapse imaging data. Built on a single-cell object-oriented framework, LivecellX integrates segmentation, tracking, feature extraction, and trajectory analysis while minimizing technical overhead and coding complexity. To address common challenges such as inaccurate segmentation, tracking errors, and lineage reconstruction, it combines automated correction with two fallback options: invalidating those single cell objects or trajectories containing errors, or manual intervention with GUI to manually correct segmentation errors or other issues that algorithms were unable to resolve. LivecellX provides a new method for resolving the segmentation and tracking errors at the same time, while the segmentation and tracking outputs can be generated with classical methods. One alternative way is performing segmentation and tracking simultaneously [25].

Leveraging machine learning, LivecellX reduces labeling burden by pretraining segmentation on public cell image datasets and employing semi-automatic algorithms for labeling incorrect segmentation and key events such as mitosis and apoptosis. LivecellX integrates transmitted-light and fluorescence features and leverages deep learning advances to extract additional information from label-free images such as organelle identification [55, 56]. In future iterations of LivecellX, we aim to incorporate more features to further enhance its functionality. Therefore, LivecellX is designed as an open, extensible framework. It supports future advances in segmentation, tracking, and integration with other analysis pipelines, enabling more robust and comprehensive studies of cellular dynamics.

## 5 Methods

### 5.1 Experimental data collection and live-cell imaging setup

#### MCF10A cells

Human breast epithelial MCF10A cells (ATCC, CRL-10317) were cultured in DMEM/F-12 medium supplemented with 5% horse serum, 20 ng/mL epidermal growth factor (EGF), 0.01 mg/mL insulin, 500 ng/mL hydrocortisone, and 100 ng/mL cholera toxin. Cells were maintained at 37 °C in a humidified incubator with 5% CO_2_ and used within 10 passages. Holo-tomographic (HT) label-free images were acquired with an Nanolive CX-A microscope (Model: 3D Cell Explorer Automated, Part Number: CX03). For each experiment, 500–5,000 cells were seeded into 35 mm glass-bottom imaging dishes (Ibidi, 80137) two days prior to imaging. Cells under the microscope were maintained at 37 °C with 5% CO_2_ using a Tokai Hit stage top incubator. HT images were acquired every 5 minutes, using a 60× objective (N.A. = 1.45). Imaging was performed over 57 hours, resulting in 684 frames. For experiments lasting over 48 hours, 300 µL of fresh medium was added every 48 hours.

#### A549 cells

Human lung adenocarcinoma A549 cells (ATCC, CCL-185) were cultured in F-12K medium supplemented with 10% fetal bovine serum (FBS) and maintained under the same incubation conditions (37 °C, 5% CO_2_). Cells within 10 passages were used. Nanolive CX-A live-cell imaging was carried out with the same setup as described above. Imaging was conducted for 27 hours at 5-minute intervals, yielding 323 frames.

#### A549/VIM-RFP cells

A549 cells stably expressing Vimentin-RFP (ATCC, CCL-185EMT) were cultured in F-12K medium (Corning) supplemented with 10% FBS. Cells were seeded into 35 mm glass-bottom culture dishes (Cellvis D35-20-1.5N) and maintained at 37 °C in a humidified atmosphere with 5% CO_2_. Cells from passages 3–10 were used for live-cell imaging with an Nikon Ti2-E microscope. For each experiment, 5,000–30,000 cells were plated two days before imaging. Differential interference contrast (DIC) images were acquired every 5 minutes, and TRITC channel images (Ex 555 nm / Em 587 nm) every 30 minutes using a 20× objective (N.A. = 0.75). Imaging was performed for 24 to 120 hours. For courses exceeding 48 hours, 500 µL of fresh culture medium was added every 48 hours.

### 5.2 Corrective Segmentation Network (CS-Net)

CS-Net is designed to refine erroneous segmentation masks by operating on cropped ROIs where single-cell segmentation errors are detected. It is implemented as a modular architecture using PyTorch Lightning and supports interchangeable backbones. In its default configuration, it employs a DeepLabV3-ResNet50 backbone pretrained on ImageNet, with the final classifier replaced by a 1 × 1 convolutional layer to produce multi-class output. An auxiliary classifier is also attached to intermediate backbone features to predict categorical segmentation error types (e.g., over-or under-segmentation).

#### Input

For each identified error region, an ROI including the erroneous mask and its surrounding context is cropped from the original image and mask. Input tensors are shaped as 3 × *H* × *W*, where *H* and *W* denote the height and width of the cropped ROI. The first channel contains the raw image. The second channel is a derived feature map (e.g., binary mask, label mask, Euclidean distance transform (EDT), or normalized EDT) as described in Section 5.2.2. The third channel duplicates the second to meet the three-channel requirement of pretrained backbones.

#### Output

CS-Net produces three output masks:

- **Segmentation mask** – A probability map representing the likelihood of each pixel belonging to a true cell object.
- **Under-seg mask** – A probability map indicating the likelihood of pixels lying outside the true object boundary but erroneously included in the original cell mask.
- **Over-seg mask** – A probability map indicating the likelihood of pixels that should be included in the cell mask but were mistakenly excluded.

#### Loss functions

The network supports cross-entropy (CE), mean squared error (MSE), and binary cross-entropy (BCE) loss functions, selected based on input types. While class and pixel-wise weighting can be applied during training, we found little performance difference with or without them. For model training, we use BCE as the loss function, specifically tailored to our requirements. Because each pixel may belong to multiple channels, it is inappropriate to use the standard cross-entropy across all classes. Instead, we compute the MSE alongside BCE, incorporating EDT to facilitate comparative evaluation.

During ablation tests, we explored various strategies for augmenting the other two channels. One approach involved removing backgrounds from raw images to create background-free representations of single cells in one of both channels. Additionally, we examined the effectiveness of incorporating segmentation masks and single-cell masks as two additional channels. This methodology enables us to assess the impact of these extra channels on the model’s performance.

#### 5.2.1 CS-Net evaluation metrics

Average Precision (AP) for object detection is typically computed using IoU-based matching; historically, *PASCAL VOC* reports AP for **bounding-box** detections at a fixed IoU threshold of 0.5 [57]. By contrast, *COCO* averages AP across multiple IoU thresholds (0.50 – 0.95 in steps of 0.05) and publishes metrics for both **bounding boxes** and **instance masks** (often denoted `AP _bbox` and `AP _mask`) [58, 59]. Bounding box-based metrics are often insufficient for detecting biologically relevant segmentation errors, as they cannot distinguish between closely packed or overlapping cells. Cell Match represents the percentage of ground-truth cells that are correctly paired with predicted cells, providing a direct measure of identification accuracy at the single-cell level. Correction success rate reports the proportion of samples in which every groundtruth cell is matched to exactly one predicted cell, serving as a stringent criterion for correctness. Intersection over Union (IoU) quantifies the overlap between predicted and ground-truth cell masks; higher IoU values indicate more accurate segmentation. Root Mean Square Error (RMSE) of normalized EDT masks measures pixel-level differences in shape between predicted and ground-truth masks, with lower values indicating greater similarity.

**Cell match rate** is defined as the percentage of predicted masks that accurately correspond to ground-truth cells. Mathematically, let *N* represent the number of samples. For the *i*-th sample, let *M*_*i*_ denote the total number of ground-truth cells, and let 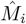 represent the number of predicted masks that correctly match a ground-truth cell based on a defined matching criterion. In our scenario, this criterion is IoU-based (IoU ≥ 0.5). Each ground-truth cell can be matched to at most one predicted cell. The cell match rate is then defined as:

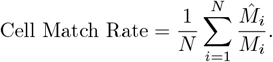

**Correction success rate** is a more stringent than the cell match rate. One segmentation error case typically relates with two or more masks. This metric requires that the predicted masks of CS-Net and ground-truth masks be *bijectively matched* within this case, meaning there must be a one-to-one correspondence between predicted masks and ground-truth masks. Each pair must satisfy a predefined matching criterion, which in our case is IoU is greater than 0.5, consistent with the criterion used in the cell match rate. A case is considered successfully corrected only if every ground-truth cell is accurately matched to a predicted cell, and vice versa. For the *i*-th case, let *G*_*i*_ represent the set of ground-truth masks, and *P*_*i*_ denote the set of predicted masks. Let ℳ_*i*_ ⊆ *G*_*i*_ *× P*_*i*_ be the set of matched pairs that satisfy the matching criterion. We define a successful correction if |ℳ_*i*_| = |*G*_*i*_| = |*P*_*i*_|, i.e., a perfect one-to-one matching exists. Then, the correction success rate is given by:

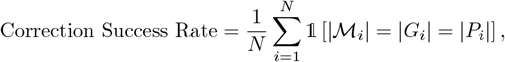

where 𝟙 [·] is the indicator function, equal to 1 if the condition is true and 0 otherwise.

**Root mean square error (RMSE)** is a metric to quantify the difference between the predicted mask and the ground-truth mask. For a predicted mask *M*_predict_ and ground-truth mask *M*_GT_, both represented as images of the same dimensions, RMSE is defined as:

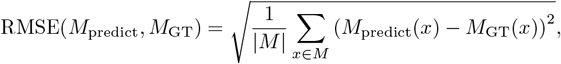

where |*M*| is the total number of pixels in the mask, and *x* indexes over all pixels. A lower RMSE signifies better alignment between predicted and ground-truth masks. RMSE is appropriate for evaluating both binary masks and EDT masks. However, when dealing with label masks, caution is advised, as the ground-truth and predicted labels may not align due to potential differences in label assignment.

#### 5.2.2 Focus mechanism

To enhance the semantic awareness of the correction task, we introduce a *focus mechanism* that guides deep learning models to concentrate on a specific cell during a single forward pass. This mechanism is implemented by incorporating a dedicated focus channel, which supplies the model with the target cell mask generated in the segmentation step. Formally, given an input image tensor *X* ∈ ℝ^*H*×*W*×*C*^ and a focus mask *F* ∈ ℝ^*H*×*W*×1^, we construct the focused input 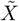 by concatenating *F* to *X* along the channel dimension:

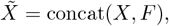

where concat(·) denotes the channel-wise concatenation operation. The resulting tensor 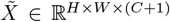 includes both the original image features and an additional semantic cue, thereby facilitating improved learning and inference.

##### Focus mask types

Different choices for *F* impact the effectiveness of the focus mechanism. We examined two primary strategies.

The first one is binary focus masks. A binary mask *F* highlights the target cell by setting the relevant pixels to 1 while marking all others as 0:

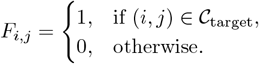

Here, 𝒞_target_ denotes the region occupied by the target cell. One issue with this method is that the focus input mask for over-seg cases may be indistinguishable from correct masks, particularly in scenarios where two over-segmented pieces of a cell are joined together. A straightforward approach is to use label masks, where each integer label represents a distinct cell, instead of binary masks. However, since these label values lack semantic meaning, they can lead to training instability. In such instances, neither the binary focus mask nor the label focus mask prove to be effective. Inspired by recent advances in the field [8, 16], we explore the potential of EDT maps as *F* to resolve these issues.

The second one is Euclidean distance transform (EDT) maps. An EDT map *F* encodes the spatial relationship between pixels and the cell boundary:

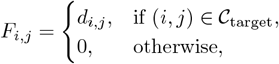

where *d*_*i*,*j*_ represents the Euclidean distance of pixel (*i, j*) to the closest boundary of the target cell. EDT maps offer a continuous gradient of focus intensity, helping to mitigate the instability issues often seen with label-based masks. Additionally, to prevent numerical and semantic problems arising from the variation in cell sizes, as illustrated in Fig. 2, we implement a normalization strategy to confine the EDT values of a cell within the range [1, range_max_]: *d*_*i*,*j*_ is normalized by first computing the scaling factor as

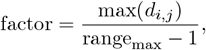

and then transforming all nonzero values (*d*_*i*,*j*_ ≥ 1) using

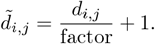

This approach ensures that the normalized EDT values are scaled relative to the maximum distance, with a minimum value of 1 for positive distances (object regions). This scaling preserves numerical stability and retains semantic meaning for both the target cell and the background regions, while also creating a numerical gap between the background and the cell boundary (Fig. 2).

#### 5.2.3 Training and testing for CS-Net

##### Data annotation

We annotated under-seg, over-seg and negative samples from the segmentation results of two models: (1) a pretrained cyto2 model from Cellpose [8] fine-tuned on our holotomography microscopy data, and (2) a custom U-Net trained on DIC images from a previous study [16]. For the Cellpose model results, we used the human-in-the-loop interface to fine-tune on annotations from 15 time frames of the A549 dataset and 35 time frames of the MCF10A dataset. The fine-tuned model was then applied to all video frames to generate segmentation masks. We created datasets via two annotation protocols: (1) manual contour drawing after visual inspection of all masks, and (2) automated computation of IoMin or IoU scores between frames *t* and *t* + 1, followed by expert annotation of triplet candidates representing possible under-seg or over-seg errors. The first method is labor-intensive but yields precise annotations, while the second enables experts to label samples with just two clicks by weakly propagating masks from neighboring time points, trading precision for efficiency.

##### Erosion and dilation

To enhance CS-Net’s robustness against segmentation errors, we apply erosion for over-seg and dilation for under-seg (Extended Data Fig.2).

The dilation and erosion of a specific pixel point in a binary mask are defined as follows:

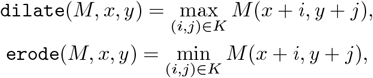

where *M* denotes a binary mask. *K* is the kernel (typically a 3 × 3 square). The functions `dilate` (*M, x, y*) and `erode` (*M, x, y*) return the dilated or eroded value of the mask *M* at pixel location (*x, y*). Dilation expands foreground regions by taking the local maximum over the kernel neighborhood, while erosion contracts them by taking the local minimum. In practice, we erode over-seg masks (*M* ← `erode` (*M*)) to emphasize disconnected boundaries and make the over-segmentation more pronounced. Conversely, we dilate under-seg masks (*M* ← `dilate` (*M*)) to further amplify the effects of merged regions. These augmentations simulate varying degrees of segmentation errors, encouraging CS-Net to generalize better. We avoid dilating over-seg masks or eroding under-seg masks, as these transformations may convert them into correct segmentations or another type of segmentation errors.

##### Varying padding sizes

Typically, an under-seg sample includes pixels from multiple adjacent cells, whereas over-seg or over-seg dropout cases may exclude parts of a cell. Since CS-Net does not have access to the true error category of an incorrect mask during inference, selecting an appropriate padding size for the cropped region of interest is essential to provide adequate information for correction.

Let *M* denote the bounding box of a single-cell mask, defined by its coordinates (*x*_min_, *y*_min_, *x*_max_, *y*_max_). We construct a padded ROI *M* ^pad^ by extending the bounding box uniformly in all directions:

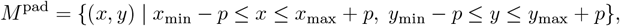

where *p* is the padding margin in pixels. To evaluate CS-Net’s robustness to contextual variation, we use multiple padding values *p* ∈ {20, 50, 80, 100} when preparing training and testing datasets. In applications, we use a fixed padding size of *p* = 50 pixels, selected based on microscope magnification and the typical pixel-scale size of individual cells.

We define *p* as a configurable hyperparameter that can be adjusted based on specific application requirements. All padding is extracted directly from the original label-free imaging data, rather than using zero-padding, to preserve spatial context and relevant background information.

##### Training data augmentation

To improve model generalization, we apply a set of spatial and photometric augmentations to both images and corresponding masks. These include random horizontal and vertical flips, affine transformations (rotation, translation, scaling, and shear), and optional Gaussian blur. All inputs are resized to 256 × 256 pixels, with bilinear interpolation for images and nearest-neighbor interpolation for masks. Augmentations are applied consistently across channels to preserve spatial alignment. The EDT channel is normalized after transformation. Normalizing images from different microscopy modalities enforces consistent intensity distributions across datasets by aligning their means and standard deviations.

##### Post-processing

For our best-performing models trained with the normalized EDT objective, we generated binary segmentation masks by thresholding the normalized EDT at values greater than 1. The resulting predicted masks were post-processed using OpenCV. Specifically, we extracted contours with `cv2.findContours`, using `mode=cv2.RETR EXTERNAL` and `method=cv2.CHAIN APPROX SIMPLE` to efficiently identify external cell boundaries.

##### Training and testing dataset composition

We assembled a comprehensive dataset for training and testing the CS-Net correction model using both holo-tomography and DIC microscopy data. The training set includes under-seg, over-seg, and negative samples, encompassing both real and synthetic errors. Specifically, HT data contributed 4,150 under-seg samples, 55 over-seg samples, 14,753 synthetic over-seg cases, 39,500 synthetic dropout-overseg samples, and 416 correct samples, with corresponding test sets comprising 1,095, 15, 3,992, 10,612, and 150 samples, respectively. The DIC training set includes diverse imaging conditions and sources, with real and synthetic under- and over-seg samples, as well as 4,084 correct examples; corresponding test sets contain 1,021 correct samples and a subset of real and synthetic segmentation errors.

### 5.3 Trajectory correction algorithms

In live-cell imaging, there are three major typical linkage distortions between single-cell masks caused by segmentation errors:

1. Under-seg of two cells from different trajectories: When two cells, one from trajectory A and the other from trajectory B, are under-segmented at time *t*, they are merged into a single mask and mistakenly assigned to the same trajectory. If the merged object is assigned to trajectory A, trajectory B typically loses a time point. In some cases, another nearby cell at time *t* may be incorrectly assigned to trajectory B instead (Fig. 3a). This type of wrong linkage can propagate, leading to a cascade of incorrect linkages during tracking, particularly those involving neighboring cells.
2. Over-seg of a single cell: If a cell at time *t* in a trajectory is segmented into multiple fragments, only one fragment is associated with the trajectory. The situation constitutes a typical over-seg case. Unlike under-seg, over-seg errors do not often result in direct tracking linkage failures. However, they can still degrade the quality of down-stream trajectory analysis. In rare instances where the segmentation model frequently produces over-seg errors, the unassigned fragments may interfere with the tracking of neighboring cells, potentially leading to incorrect trajectory linkages (Fig. 3b).
3. Absent cell: When the segmentation algorithm fails to detect an existing cell at time *t*, no mask is produced for that location. As a result, the tracking algorithm may terminate the trajectory prematurely or mis-assign the cell in subsequent frames. This type of error can also cause confusion in nearby trajectories if the algorithm incorrectly links neighboring cells to compensate for the missing one (Fig. 3c).

To efficiently address tracking and segmentation errors in large-scale live-cell imaging, we developed an iterative trajectory correction algorithm that integrates spatial-temporal consistency checks with CS-Net (Fig. 3e). The method is parameterized by values including a temporal search window (3 frames), correction crop padding (20 pixels), a maximum number of correction iterations (default 100), and an exclusion zone for boundary cells (50 pixels). All parameters are tunable.

For under-seg, the algorithm inspects each trajectory for missing frames. When a gap is found, it examines whether any cell masks present in the missing frame overlap substantially with the trajectory’s masks in adjacent frames, using an IoMin/IoU threshold=0.5. If such matches exist, the candidate error region is cropped and passed to the trained CS-Net, which classifies and corrects the segmentation. The resulting corrected masks are assigned back to the trajectories using the linear assignment problem (LAP) based on spatial relation. If the correction is successful, the trajectories are updated accordingly: splitting, merging, or extending as needed.

For over-seg, the criterion we used in the reported analyses is based on morphological dynamics. Here, the algorithm monitors the area of each segmented cell mask over time and flag instances where the area changes abruptly beyond a predefined threshold. If such a candidate is detected, the local region is again input to CS-Net for correction and classification. For over-seg, typically there is only one trajectory involved, so the corrected single-cell mask is inserted directly back into the trajectory without any assignment step.

The entire process is repeated iteratively. After each global correction iteration, the trajectories are updated, and new candidate errors are detected based on the most recent corrections. Iterations continue until no further correction is possible or the maximum number of rounds is reached. In all cases, if a correction cannot be confidently corrected, the related linkages are left unchanged and the error is flagged, but not retried unless its context is modified in subsequent rounds. Users can utilize LivecellX GUI to visually inspect these samples and perform manual correction.

In tracking algorithm like SORT tracker [47], the maximum age parameter (max gap in frames) defines the largest number of consecutive frames a trajectory can persist without a successful linkage. Hence the impact of this parameter on the correction process should be evaluated.

The number of over-seg cases in our HT microscopy dataset is significantly low compared to under-seg cases. To address this imbalance, we synthesized 5,000 over-seg samples using the same protocol employed for generating synthetic over-seg training data. We evaluated our trajectory correction algorithms on both the synthetic and real over-seg datasets.

#### 5.3.1 Trajectory correction evaluation metrics

We introduce **trajectory vacancy** as a straightforward yet informative metric for evaluating the quality of a single-cell trajectory. For a given trajectory, the vacancy rate quantifies the extent to which it is fragmented over time. Specifically, we define the trajectory vacancy rate as follows::

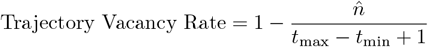

where *t*_max_ and *t*_min_ are the maximum and minimum time frames of the cells in the trajectory after tracking, and 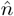 is the number of tracked cells in the trajectory. A lower vacancy rate indicates a more complete and continuous trajectory.

### 5.4 Biological process detection and lineage reconstruction

LivecellX supports automated classification of single-cell trajectories derived from temporal features and time-lapse image sequences. Let each single cell trajectory be represented as a fixed-length sequence of feature vectors:

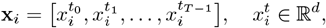

where *t*_0_ is centered around the cell’s birth event and *T* (typically ≈ 8) defines the length of the sliding window sampling temporal context. Each trajectory **x**_*i*_ belongs to a biological process label *y*_*i*_ ∈ {1, …, *C*}, indicating its class over the entire sequence. The goal is to train a model *f*_*θ*_ that maps whole trajectories to their corresponding classes:

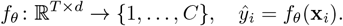

This formulation ensures classification captures dynamic cellular behaviors rather than static snapshots.

LivecellX facilitates the preparation of machine learning–ready inputs by providing helper functions that format temporal features, image patches, and video sequences directly from its data structures (e.g., single-cell and trajectory objects). The modular detection pipeline is extensible, enabling users to plug in custom classifiers *f*_*θ*_. We provides pretrained CNN- and transformer-based models optimized for our DIC and HT A549 datasets produced in our study.

Lineage reconstruction in LivecellX is carried out after detecting biological processes. The framework provides utility functions to split or merge trajectories based on common cellular events, such as mitosis and cell fusion. In the case of mitosis, the mother and daughter trajectories are separated at the time point of division. For cell fusion, two individual cell trajectories are combined into a single trajectory at the fusion time point. The relationships between cell trajectories are stored within the LivecellX data structures. Users have the flexibility to define new types of relationships either programmatically or through the graphical user interface.

To facilitate lineage visualization, LivecellX provides a light-weight tree traversal utility based on depth-first search (DFS). Within this algorithm, each trajectory is represented as a node in the lineage tree, and mother-daughter relationships are managed via the trajectory object structure. Starting from a root cell trajectory, the DFS-based algorithm recursively visits descendants, generating ordered lineage sequences or visual renderings.

### 5.5 Feature extraction and trajectory dynamics

The module performs feature extraction using the active shape model, Haralick features, local binary pattern, variational auto-encoder, Harmony [51], and diffusion models [60, 61]. LivecellX integrates the active shape model implemented in Celltool [32]. Haralick features and local binary patterns are extracted using the Mahotas package [62]. The variational auto-encoder is implemented through Harmony. LivecellX provides helper functions for constructing time delay embeddings from trajectories.

#### Harmony VAE for Microscopy Image Disentanglement

We adopted the original Harmony framework [51], a generic VAE-based model designed to disentangle semantic content from parameterized transformations in an unsupervised manner. Harmony combines a standard encoder–decoder architecture with a cross-contrastive learning objective to separate transformation-specific and content-specific latent factors. In its original form, Harmony demonstrated strong performance on structured datasets such as MNIST and cryo-EM images. However, we found that this architecture does not generalize effectively for our label-free microscopy datasets, where high noise levels and fine-scale cellular heterogeneity pose additional challenges. A primary limitation of the original model in our setting was posterior collapse, where the learned latent variables failed to capture meaningful variation and reconstructions became nearly identical regardless of latent input [8]. Additionally, the model often produced nearly identical reconstructions across different latent codes, indicating poor disentanglement and underfitting of the semantic space. To address these shortcomings, we replaced Harmony’s original fully connected encoder and decoder with convolutional architectures tailored for high-resolution microscopy images. Specifically, we employed a deeper convolutional encoder with residual blocks and batch normalization to extract hierarchical visual features, and a mirrored transposed convolutional decoder to enhance spatial resolution during reconstruction. These architectural changes substantially increased the model capacity and improved its ability to capture both global cell shape and fine-grained morphological variations. We retained Harmony’s cross-contrastive loss structure, which includes three dissimilarity terms—between the image reconstructed from semantic latent space and the image transformed by the predicted transformation parameters, between the transformed outputs of original and augmented views, and between the reconstruction from the augmented input and its transformed version—along with a KL divergence term enforcing proximity of semantic latent distributions between similar inputs. These components jointly drive the disentanglement of latent representations while avoiding trivial solutions. In our implementation, we further regularized transformation parameters through appropriate activation functions and shared kernel weights. We empirically found that our convolutional Harmony variant better disentangles latent content from cell shape variability and imaging transformations. This enhanced formulation enables generation of synthetic microscopy images and improves robustness to transformation-induced noise.

#### Harmony VAE: Original Formulation and Our Adaptations for Label-free Microscopy Images

The Harmony model aims to disentangle semantic content *z* ∈ ℝ^*d*^ from transformation parameters *k* ∈ ℝ^*m*^ using a variational autoencoder framework combined with a contrastive learning objective.

Let *x* ∈ 𝒳 be an input image. The encoder maps *x* to both transformation parameters and a variational distribution over semantic latent variables:

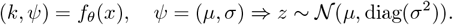

The decoder reconstructs from the latent code, while the transformation module synthesizes a transformed image:

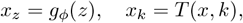

where *T* (·, *k*) denotes a differentiable transformation function parameterized by *k*.

Harmony’s training objective is defined as:

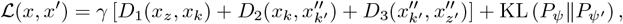

where *x*^*′*^ = *T* (*x, k*^*′*^) is a randomly transformed view, and 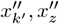 are outputs from applying the encoder and decoder to *x*^*′*^. Each *D*_*i*_ denotes a dissimilarity function, typically mean squared error (MSE).

#### Modifications on Harmony

To adapt Harmony for high-resolution and morphologically complex microscopy data, we replaced the fully connected encoder–decoder architecture with CNNs to better capture spatial structure.

##### Modified Encoder

Instead of the original MLP-based encoder,

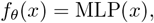

we use a convolutional encoder:

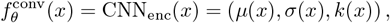

where *µ*(*x*) and *σ*(*x*) are obtained from global average pooled convolutional features followed by linear projections.

##### Modified Decoder

Replacing the MLP decoder,

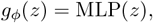

we use a convolutional decoder:

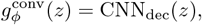

consisting of upsampling and convolutional blocks to reconstruct high-resolution images.

##### Updated Forward Pass

The modified reconstruction path becomes:

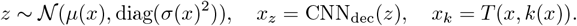

This convolutional formulation enhances Harmony’s representational capacity, improves robustness to complex transformations, and effectively mitigates posterior collapse in microscopy imaging applications.

## 6 Data Availability

Raw live-cell label-free images are stored and shared on OSF.

## 7 Code Availability

The `LivecellX` package is publicly available at: https://github.com/xing-lab-pitt/livecellx. The repository includes source code, installation instructions, and example notebooks to facilitate reproduction and further development.

## Acknowledgements

This work was partially supported by National Science Foundation (MCB2205148) to JX, National Institute of Biomedical Imaging and Bioengineering (T32EB009403) to DP, National Natural Science Foundation of China Grants (No.12247104) to WW and the University of Pittsburgh Center for Research Computing and Data, `SCR_022735` through the resources provided. Specifically, this work used the H2P cluster, which is supported by NSF award number (OAC-2117681).

## Declarations

The authors declare no conflict of interest.

## Notes

### Competing Interest Statement

The authors have declared no competing interest.

### Summary of Updates

Figure1-6 are updated and the manuscript are reviesed

