## supplement for "LivecellX: Corrective Deep Learning for Object-Oriented Single-Cell Analysis in Live-Cell Imaging"

### Supplementary Figures

October 30, 2025

#### Extended Data Figures

a

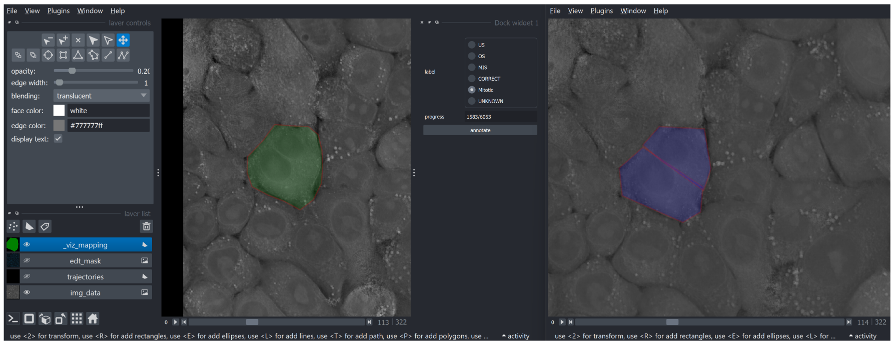

b

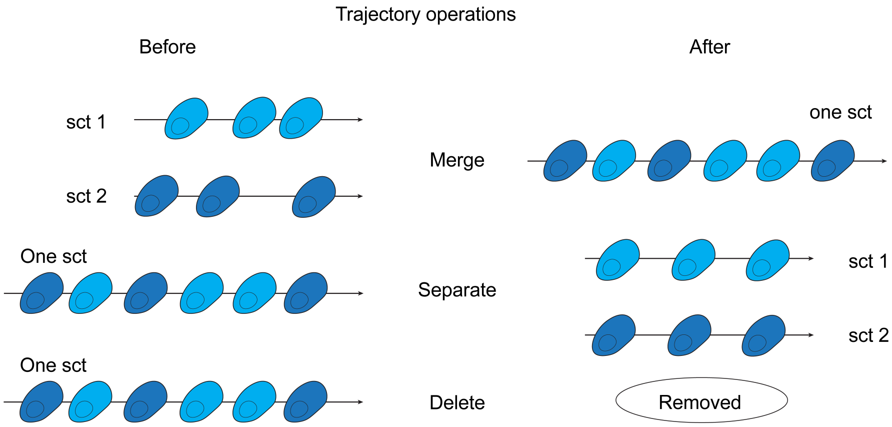


**Extended Data Figure 1. Segmentation error propagation and annotation workflow in LivecellX. a**, Graphical user interface (GUI) for manual annotation in LivecellX. **b**, Types of trajectory operations in LivecellX.

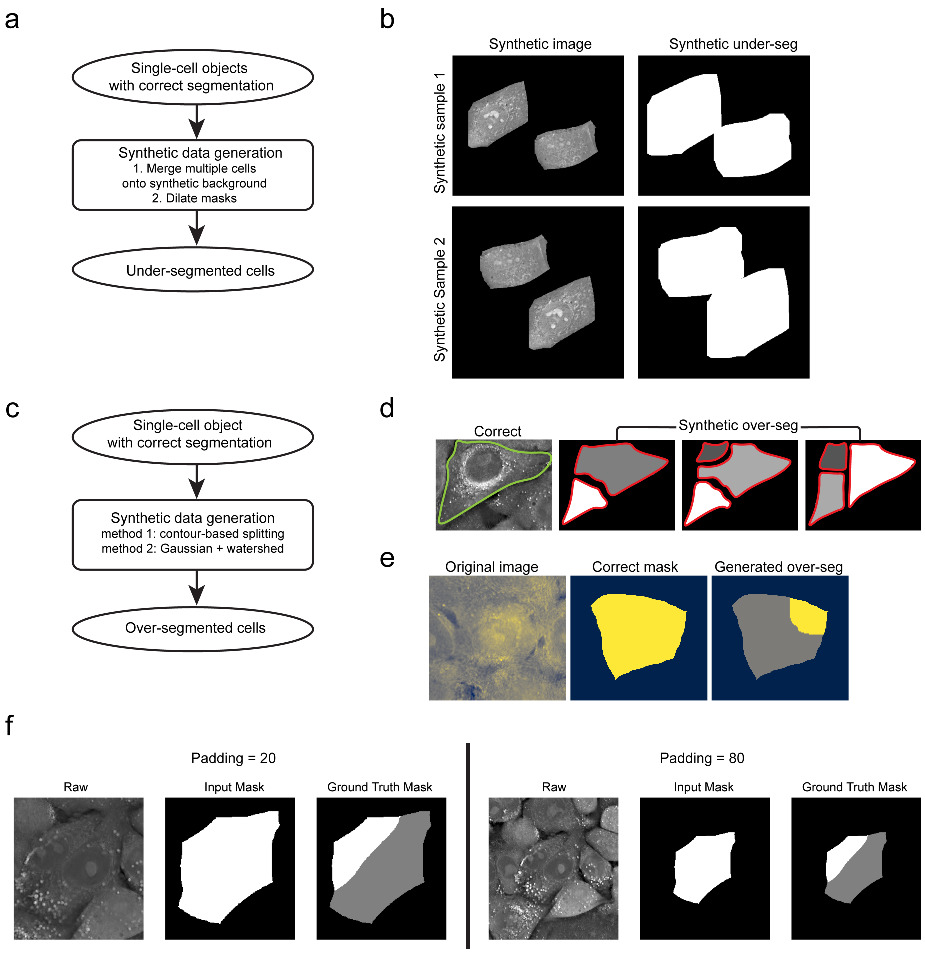


**Extended Data Figure 2. Synthetic generation of under-seg and over-seg training data. a**, Workflow for generating synthetic under-seg samples. Multiple single-cell objects with correct masks are merged onto a synthetic background to create under-segmented samples. **b**, Typical synthetic under-seg examples. For each synthetic sample (rows), left panel shows the composite synthetic image, and right panel shows the corresponding merged mask where multiple cells are fused into a single under-segmented region. **c**, Workflow for generating synthetic over-seg samples. Single-cell objects with correct segmentation are processed using two synthetic data generation strategies: (1) contour-based splitting, and (2) randomly selecting internal seed points and adding Gaussian intensity peaks, followed by watershed segmentation to divide the cell mask. **d**, Typical synthetic over-seg examples. A correct mask (left) is split into multiple fragments (middle and right) to simulate over-segmentation, with ground-truth boundaries (green) and predicted over-seg boundaries (red) highlighted. **e**, Example of an original image (left), corresponding correct mask (middle), and generated over-seg mask (right) using the synthetic over-seg sample generation pipeline. **f**, A typical training sample with different number of padding pixels.

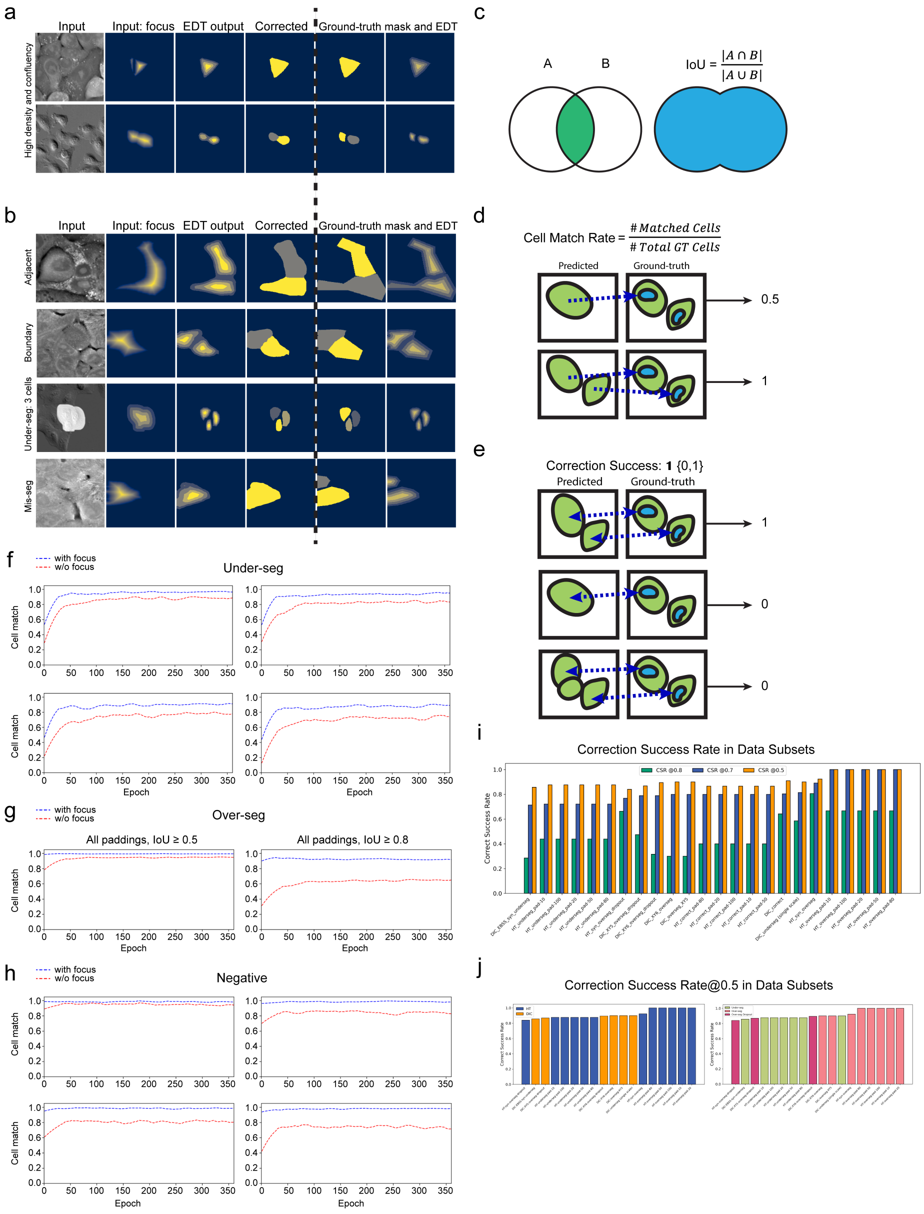


**Extended Data Figure 3. Evaluation metrics and case studies for CS-Net segmentation correction.** **a**, Representative examples of CS-Net correction on high-density/confluent samples. Each row is shown with (from left to right): input image, input mask with EDT focus channel, CS-Net output, corrected segmentation, and ground-truth mask with corresponding EDT. **b**, Additional challenging cases including adjacent-cell merges, ambiguous boundaries, under-seg involving three cells, and un-categorized mis-segmentation. The columns follow the same layout as in (a). **c**, Schematic of intersection over union (IoU) metric as used for mask comparison, defined as the ratio of the intersection area to the union area between predicted (A) and ground-truth (B) masks. **d**, Illustration of the cell match rate metric, defined as the proportion of correctly matched cells between prediction and ground truth; examples shown represent cell match rate equals fifty percent (top) and one hundred percent(bottom). **e**, Schematic for the correction success rate metric, indicating cases where the predicted mask set achieves perfect bijective correspondence with the ground truth. **f**–**h**, Learning curves of CS-Net cell match rate (y-axis) versus training epoch (x-axis) under different padding settings and error types. **f**, under-seg cases; **g**, over-seg cases; **h**, negative (correct) cases. Each subplot compares different padding sizes(Top left:20; Top right: 50; Bottom left: 80; Bottom right: 100) in **f** and **h**. And plots in **g** show performance for IoU $\geq 0.5$ (dashed blue) and $\geq 0.8$ (dotted red) trained using data augmented with all padding sizes. **i**, Bar plots showing the correction success rates across different data subsets at three thresholds (IoU $\geq 0.5$, IoU $\geq 0.7$, IoU $\geq 0.9$), highlighting variation in performance depending on error type and padding configuration. **j**, Correction success rates at IoU $\geq 0.5$ for each data subset, displayed separately for different microscopy modalities (left) and error-type (right) splits, enabling comparison of generalization performance across conditions.

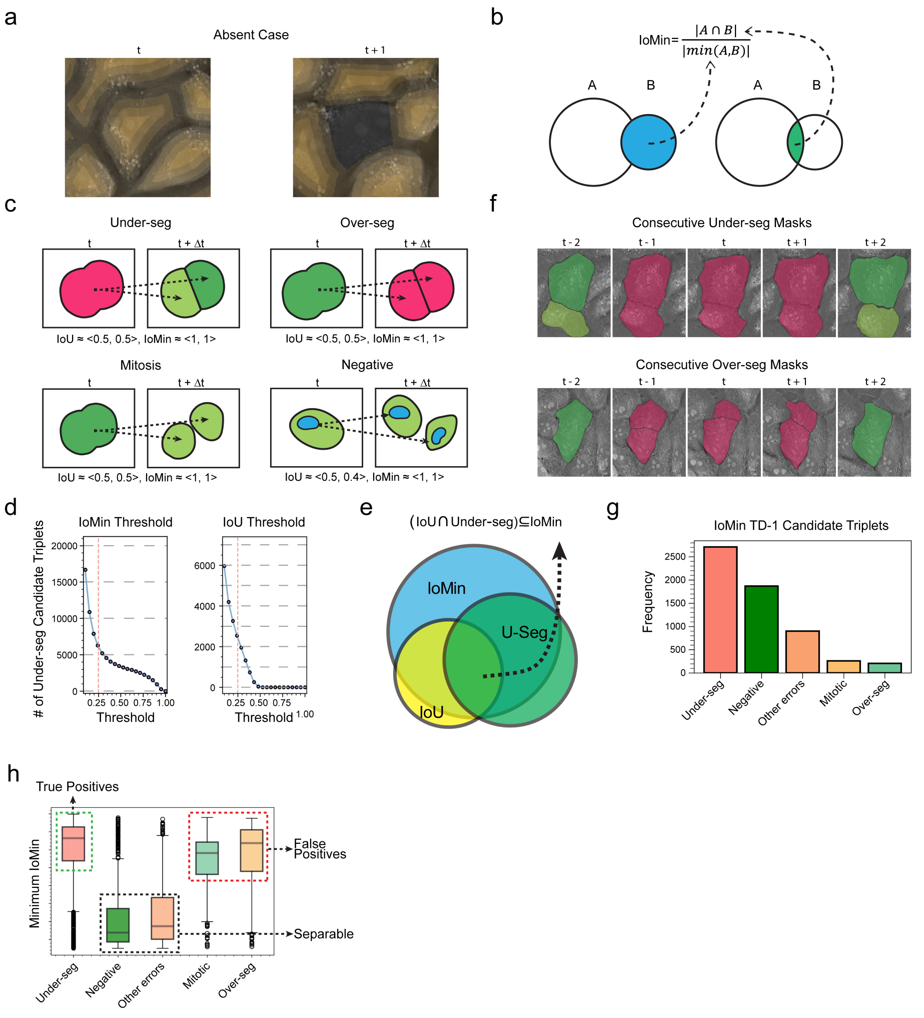


**Extended Data Figure 4. Temporal inconsistency detection and analysis of segmentation errors using IoMin and IoU criteria.** **a**, Example of an absent cell between consecutive frames. **b**, Definition and schematic illustration of the Intersection-over-Minimum (IoMin) metric. **c**, Schematic representation of candidate triplet configurations (under-seg, over-seg,mitosis and negative(no error)) between consecutive frames for error detection. Metric values for IoU and IoMin annotated. **d**, Threshold analyses for IoMin and IoU, showing the number of under-seg candidate triplets detected as a function of each metric's threshold. **e**, Venn diagram illustrating the relationship between IoMin, IoU, and under-seg detection criteria in the HT testing dataset,showing that the intersection of IoU and true under-seg cases forms a subset of IoMin. **f**, Example of consecutive under-seg masks (top)and over-seg masks(bottom) over time, demonstrating persistent segmentation error propagation across frames. **g**,

Frequencies by error type in TD-1 candidate triplet obtained with IoMin metric demonstrating the predominance of under-seg cases among detected candidates. **h**, Boxplots showing the distribution of minimum IoMin values for each error class, illustrating separability and overlap between true positives and false positives.

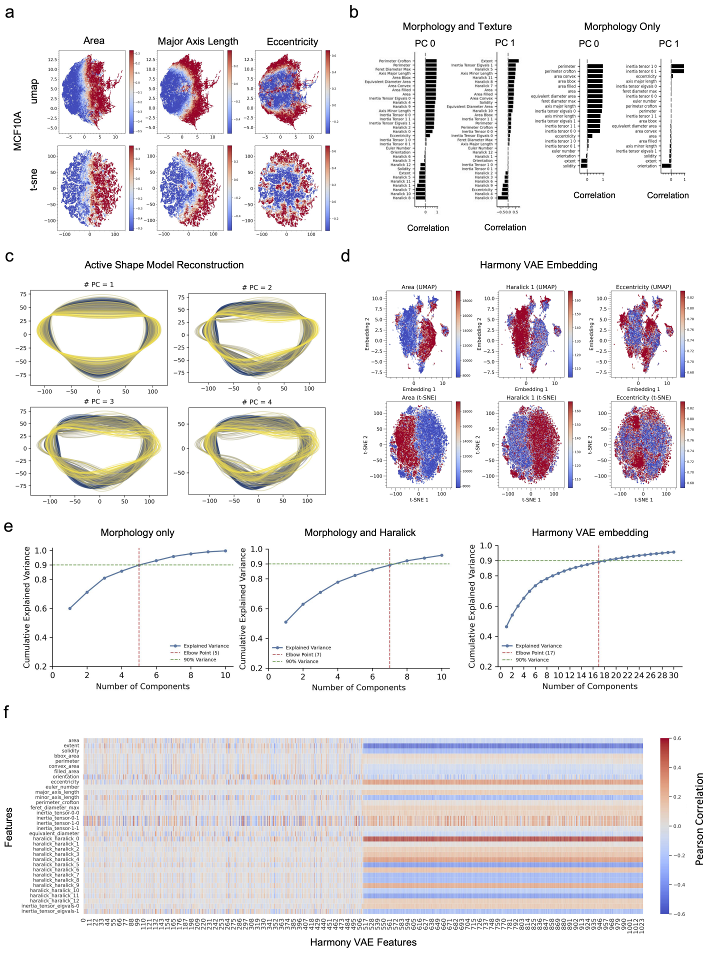


**Extended Data Figure 5. Dimensionality reduction, feature embedding, and cross-modal comparison of cell morphological features. a**, Visualization of key morphological features (area, major axis length, eccentricity) for MCF10A cells using UMAP and t-SNE. **b**, Correlation between morphology and text features with the principal components (PC0 and PC1) of single cells represented with both morphology and text features or morphology features only, indicating the most discriminative features in each component. **c**, Reconstructions of single-cell contours in active shape model with different numbers of PCs. **d**, Visualizing comparison of Harmony VAE embeddings with key morphology and texture features with UMAP and t-SNE. Colors represent feature values. The correspondence between clusters and colors reflect correlation between them. **e**, Cumulative explained variance curves for dimensionality reduction of (left) morphology-only features, (middle) morphology combined with Haralick texture features, and (right) Harmony VAE embeddings. The intersections of red and green dashed lines indicate the number of components required to capture 90% of total variance. **f**, Heatmap showing Pearson correlation of morphology and texture features with Harmony VAE embedding features, highlighting concordance and complementarity between learned and traditional features.

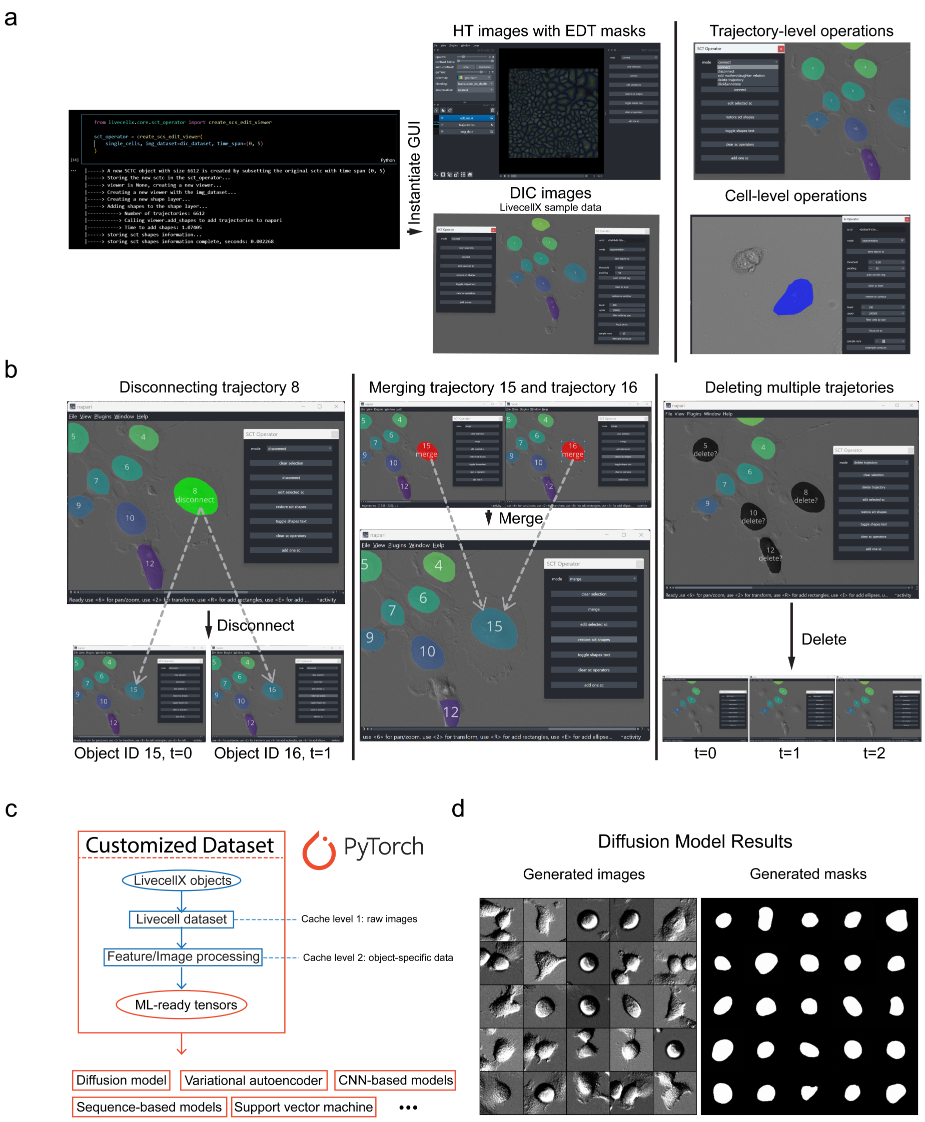


**Extended Data Figure 6. LivecellX GUI functionality and generative diffusion model results.** **a**, Overview of the LivecellX Napari-based GUI. Shown are code to instantiate the GUI (left), examples of HT images with EDT masks (top middle), DIC images from LivecellX sample dataset(bottom middle), available operations at both the trajectory drop-down menu (top right) and single-cell operations (bottom right). **b**, Demonstration of key interactive operations within the GUI, including disconnecting specific trajectories(left), merging selected trajectories(middle), and deleting multiple trajectories with one click (right). **c**, Integration of LivecellX with PyTorch for scalable machine learning workflows. The LivecellX framework enables conversion of object-level data from the Livecell data structures into machine learning–ready tensors. This pipeline includes raw image caching (cache level 1), feature and image processing for object-level representation (cache level 2), and direct compatibility with PyTorch. The resulting dataset can be used to train and evaluate a wide variety of models including diffusion models, variational autoencoders, CNNs, sequence-based models, and statistical methods like SVMs, supporting advanced analyses of single-cell dynamics. **d**, Results from a generative diffusion model trained on LivecellX data objects constructed from DIC images. Left: representative examples of generated single-cell images. Right: corresponding generated cell-body segmentation masks, demonstrating the model's ability to capture realistic cell morphology and mask variability.
